# Interference with Systemic Negative Feedback Regulation as a Potential Mechanism for Nonmonotonic Dose-Responses of Endocrine-Disrupting Chemicals

**DOI:** 10.1101/2024.09.04.611257

**Authors:** Zhenzhen Shi, Shuo Xiao, Qiang Zhang

## Abstract

**Background:** Endocrine-disrupting chemicals (EDCs) often exhibit nonmonotonic doseresponse (NMDR) relationships, posing significant challenges to health risk assessment and regulations. Several molecular mechanisms operating locally in cells have been proposed, including opposing actions via different receptors, mixed-ligand heterodimer formation, and receptor downregulation. Systemic negative feedback regulation of hormone homeostasis, which is a common feature of many endocrine systems, has also been invoked as a mechanism; however, whether and how exactly such global feedback structure may underpin NMDRs is poorly understood.

**Objectives:** We hypothesize that an EDC may compete with the endogenous hormone for receptors (i) at the central site to interfere with the feedback regulation thus altering the physiological hormone level, and (ii) at the peripheral site to disrupt the hormone action; this dual-action may oppose each other, producing nonmonotonic endocrine effects. The objective here is to explore – through computational modeling – how NMDRs may arise through this potential mechanism and the relevant biological variabilities that enable susceptibility to nonmonotonic effects.

**Methods:** We constructed a dynamical model of a generic hypothalamic-pituitary-endocrine (HPE) axis with negative feedback regulation between a pituitary hormone and a terminal effector hormone (EH). The effects of model parameters, including receptor binding affinities and efficacies, on NMDR were examined for EDC agonists and antagonists. Monte Carlo human population simulations were then conducted to systemically explore biological parameter conditions that engender NMDR.

**Results:** When an EDC interferes sufficiently with the central feedback action of EH, the net endocrine effect at the peripheral target site can be opposite to what is expected of an agonist or antagonist at low concentrations. J/U or Bell-shaped NMDRs arise when the EDC has differential binding affinities and/or efficacies, relative to EH, for the peripheral and central receptors. Quantitative relationships between these biological variabilities and associated distributions were discovered, which can distinguish J/U and Bell-shaped NMDRs from monotonic responses.

**Conclusions:** The ubiquitous negative feedback regulation in endocrine systems can act as a universal mechanism for counterintuitive and nonmonotonic effects of EDCs. Depending on key receptor kinetic and signaling properties of EDCs and endogenous hormones, some individuals may be more susceptible to these complex endocrine effects.

## Introduction

Endocrine-disrupting chemicals (EDCs) are diverse groups of compounds that interfere with the production, metabolism, transportation, and actions of endogenous hormones. The disrupting effects can be mediated through a variety of mechanisms, including perturbation of hormone synthesis, dysregulation of metabolic enzymes, competition for plasma binding proteins, and acting as hormone receptor agonists or antagonists (Combarnous and Nguyen 2019, La Merrill, Vandenberg et al. 2020). Broadly studied EDC families include polychlorinated biphenyls, polybrominated biphenyls, dioxins, bisphenol A (BPA), dichlorodiphenyltrichloroethane, and many pharmaceutical compounds (Chen, Yang et al. 2022). Numerous animal studies have demonstrated that by disrupting the various endocrine systems, these EDCs can produce a plethora of adverse health outcomes, including defects in development, reproduction, metabolism, and immunity, and cancer (Diamanti-Kandarakis, Bourguignon et al. 2009, Boas, Feldt-Rasmussen et al. 2012, Sifakis, Androutsopoulos et al. 2017, Ghassabian and Trasande 2018). Emerging epidemiological studies also reveal that many human health disorders are associated with exposures to environmental EDCs, even at low exposure levels (Skakkebaek, Rajpert-De Meyts et al. 2001, Delbès, Levacher et al. 2006, Hatch, Troisi et al. 2006, Alonso-Magdalena, Quesada et al. 2011, Wan, Co et al. 2022, Szczęsna, Wieczorek et al. 2023). Thus, the human health risk of environmental EDCs is a significant public health concern.

One of the main challenges in assessing the health risks of EDCs is that it is not straightforward to translate the toxicities observed in high-dose animal studies into health outcome predictions for environmentally relevant low-dose exposures in humans. A major issue here is that EDCs have been widely reported to exhibit nonmonotonic dose-response (NMDR) behaviors, where their biological effects can change directions in a dose-dependent manner, presenting as J/U or Bell (inverted U) shapes (Vandenberg, Colborn et al. 2012, Lagarde, Beausoleil et al. 2015, Soto and Sonnenschein 2024). NMDRs have been observed *in vitro* as well as *in vivo* on multiple biological endpoints including organ weights, uterine growth, mammary gland development, immune response, and hormone levels for a variety of EDCs (Program 2001, Bloomquist, Barlow et al. 2002, Ahn, Hu et al. 2005, Narita, Goldblum et al. 2006, Shioda, Chesnes et al. 2006, Wadia, Vandenberg et al. 2007, Dickerson, Guevara et al. 2009, Cabaton, Wadia et al. 2010, Lagarde, Beausoleil et al. 2015, Badding, Barraj et al. 2019, Montévil, Acevedo et al. 2020). NMDRs have also been reported in many epidemiological studies between environmental EDCs and a variety of health endpoints (Vandenberg, Colborn et al. 2012). For instance, a U-shaped curve was reported for the relationship between serum PCB178 and HDL (Lee, Steffes et al. 2011), Bell-shaped curves were reported for the relationships between the serum polybrominated diphenyl ether 153 and diabetes/metabolic syndrome risks and triglyceride levels (Lim, Lee et al. 2008), between serum PCB congeners or organochlorine pesticides and BMI, plasma lipid, and insulin resistance (Lee, Steffes et al. 2011), and lately between serum total effective xenoestrogen burden level and endometrial cancer risk (Costas, Frias-Gomez et al. 2024).

When a chemical displays NMDR, the linear and linear non-threshold extrapolation methods as well as the Benchmark Dose (BMD) modeling approach can no longer be applied to estimate the low-dose risk or reference dose in the framework of traditional risk assessment, thus posing regulatory challenges (EFSA Scientific Committee, More et al. 2021). Following a couple of Scientific and Position Statements on EDCs in the past decade (Diamanti-Kandarakis, Bourguignon et al. 2009, Gore, Chappell et al. 2015), The Endocrine Society has recently gone as far as stating that “Regulatory toxicology should implement endocrine concepts such as low dose and NMDR without further delay. Because of the presence of NMDR, it cannot be assumed that there are thresholds below which EDC exposures are safe” (The Endocrine Society 2018).

While observational studies have provided much of the evidence that NMDRs are not uncommon with EDCs and have caused concerns in the endocrine community, some doubts still remain as to whether NMDRs are bona fide biological phenomena or just experimental artifacts that have escaped alternative interpretations (Heindel, Newbold et al. 2015, Camacho, Lewis et al. 2019). The field has reached a point that conducting more observational studies is of limited value; rather, more mechanistic investigations, theoretical or experimental, need to be pursued to better understand the operation of the hormone signaling pathways and the physiological conditions under which nonlinear endocrine effects may arise (Birnbaum 2012). A number of biological mechanisms have been postulated for the NMDR behaviors of EDCs. Notwithstanding cytotoxicity, these potential mechanisms include (i) divergent biological actions via two distinct nuclear receptors, (ii) incoherent feedforward through membrane and nuclear receptors, (iii) ligand-induced receptor desensitization or degradation, (iv) divergent effects of the parent compound and its metabolite, (v) coactivator squelching, (vi) induction of repressor, and (vii) negative feedback regulation (Kohn and Portier 1993, Kohn and Melnick 2002, Conolly and Lutz 2004, Li, Andersen et al. 2007, Vandenberg, Colborn et al. 2012, Cookman and Belcher 2014, Lagarde, Beausoleil et al. 2015, Xu, Liu et al. 2017).

Experimental validation of these proposed mechanisms is rare (Villar-Pazos, Martinez-Pinna et al. 2017). In contrast, a number of computational studies have investigated several NMDR mechanisms (Kohn and Portier 1993, Kohn and Melnick 2002, Conolly and Lutz 2004, Li, Andersen et al. 2007). These models examined the NMDR effects within the classical framework of nuclear receptor-mediated endocrine signaling in cells targeted by EDCs. Kohn and Melnick showed that when agonist-bound receptors recruit coactivators with lower affinity than the endogenous hormone-bound receptors, as the agonist increases in concentration to replace hormone-bound receptors, the induced gene expression will eventually reverse direction and decrease, producing an inverted U-shaped NMDR (Kohn and Melnick 2002). Conolly and Lutz demonstrated that for an agonist X, when a mixed-ligand heterodimer is formed between the endogenous hormone-liganded receptor monomer and X-liganded receptor monomer, a U-shaped NMDR can arise if the heterodimer is transcriptionally inactive (Conolly and Lutz 2004). The model was used to explain the U-shaped response observed with flutamide in androgen receptor reporter assays (Maness, McDonnell et al. 1998), and the existence of mixed-ligand heterodimers has been experimentally demonstrated (Leonhardt, Altmann et al. 1998). We further proposed that because receptor homodimerization is an inherently nonlinear mass-action process, U-shaped NMDR can arise for an agonist even in the absence of the mixed-ligand heterodimer (Li, Andersen et al. 2007). Our mathematical model further extended that this U-shaped dose response can be enhanced if mixed-ligand heterodimers can also be formed and are transcriptional repressors. Alternatively, if the mixed-ligand heterodimers are transcriptional activators, inverted U-shaped NMDR can arise. In summary, these mathematical modeling studies provided valuable insights into the local molecular mechanisms of NMDR in cells of target tissues.

The molecular events associated with these local NMDR mechanisms described above are not necessarily unique to hormonal signaling in endocrine systems. The fact that EDCs are more frequently observed to produce NMDR than non-EDCs suggests that some common features of the endocrine systems may be the most likely underlying mechanism. A prominent feature of an endocrine system is that the participating tissue/organ components are organized in a systemic negative feedback loop to maintain hormone homeostasis, as exemplified by the hypothalamic-pituitary-endocrine (HPE) axes for many hormones. For instance, in the hypothalamic-pituitary-thyroid (HPT) axis, the negative feedback inhibition of TRH and TSH by T4 and T3 is essential to maintain the circulating thyroid hormone levels within a narrow range (Costa-e-Sousa and Hollenberg 2012). Such global negative feedback regulation has been suggested as a potential mechanism underpinning some NMDR phenomena (Vandenberg, Colborn et al. 2012, Lagarde, Beausoleil et al. 2015). It has been argued that negative feedback can result in temporal nonmonotonic fluctuations in hormone levels, i.e., transient rise and fall over time in response to acute perturbations by EDCs (Vandenberg, Colborn et al. 2012, Lagarde, Beausoleil et al. 2015). For chronic environmental exposures, steady-state NMDR is particularly important. However, it is not known whether negative feedback is capable of producing NMDR at steady-state conditions, and if so, how and under what biological conditions it may occur.

By competing with the endogenous hormone for cognate receptors, an EDC agonist or antagonist can hit an endocrine system at two separate sites – one is the peripheral target site where the endogenous hormone exerts its biological effects, and the other is the central feedback site, such as the brain in an HPE axis, where the synthesis and release of a pituitary hormone (and the corresponding hypothalamic releasing hormone) is regulated by the endogenous hormone. We hypothesize that when an EDC has differential signaling strengths thus actions in the two sites, an NMDR may arise. In the present study, we constructed a minimal mathematical model of a generic HPE feedback system to investigate this hypothesis. We discovered that the binding affinities and efficacies of the EDC for the central and peripheral receptors, relative to those of the endogenous hormone, follow a set of specific rules to enable J/U or Bell-shaped NMDRs. We then extended the HPE model to a virtual human population model to further explore the biological variabilities that may make some individuals susceptible to NMDR outcomes.

## Methods

### 1. Construction of a minimal mathematical model of HPE feedback loop

Here we used the generic HPE feedback framework to build a minimal mathematical model to investigate the biological conditions for emergence of NMDRs. Other endocrine feedback regulation systems not involving the hypothalamus and pituitary, such as those between insulin and glucose, and between parathyroid hormone, vitamin D3 (VD3), and calcium, should work in a similar fashion. The structure of the dynamic model of the HPE feedback loop is illustrated in Fig. 1A. It consists of peripheral and central modules. The interactions between the hormones, receptors, and EDCs are modeled based on the law of mass action. In the peripheral module, the production of the effector hormone (*EH*) is stimulated by the pituitary hormone (*PH*) in a first-order manner with a rate constant *k*_1_. *EH* is degraded in a first-order manner with a rate constant *k*_2_. *EH* binds reversibly to the peripheral receptor (*PR*) to form a ligand-receptor complex *EHPR* in a target tissue, with a second-order association rate constant *k*_5*f*_ and first-order dissociation rate constant *k*_5*b*_. *EHPR* produces an Endocrine Effect (*EE*) proportional to its concentration.

**Figure 1.**
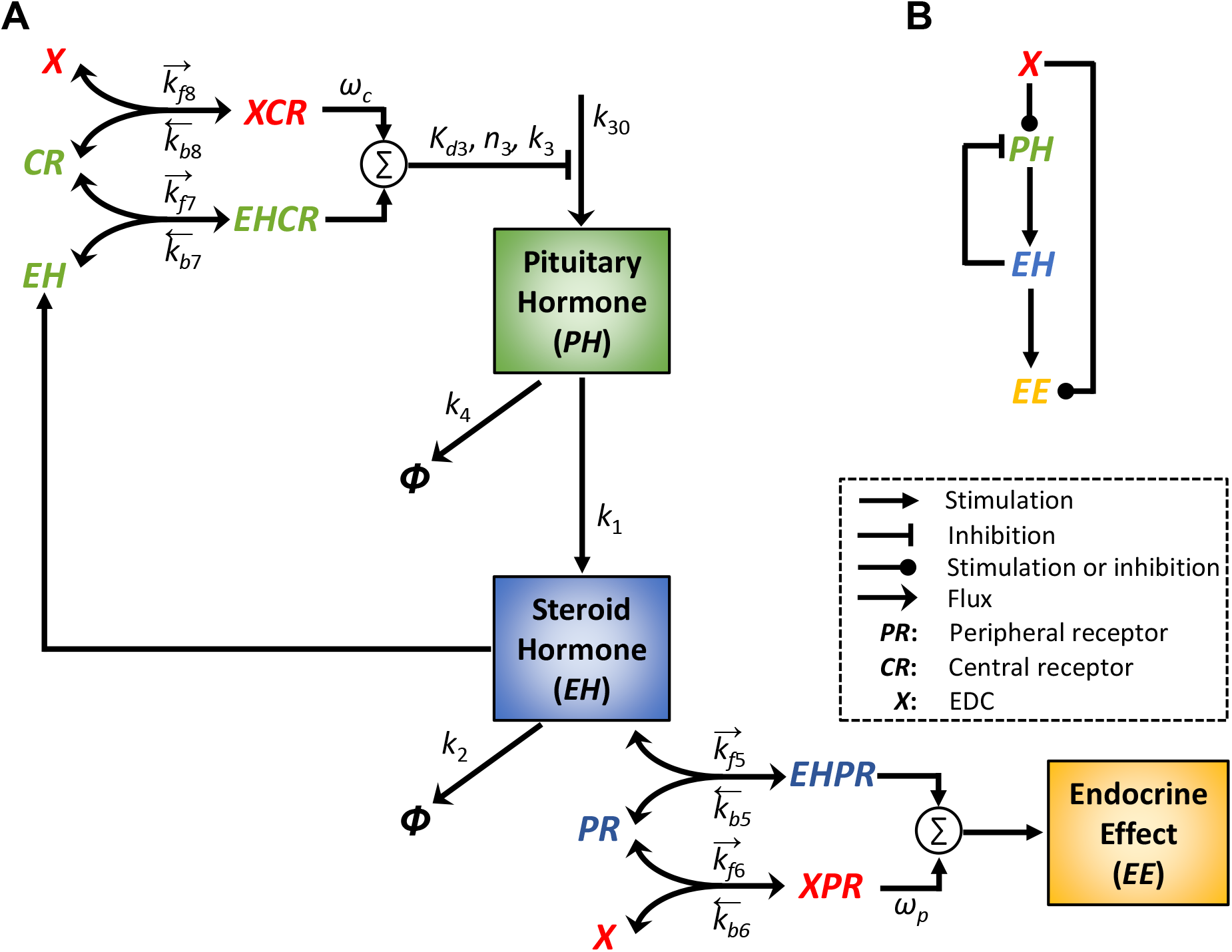
Schematic diagram of the minimal HPE feedback model and its perturbation by an EDC *X*. **(A)** Details of the model structure with parameters indicated (see Methods for details). Arrows on the top of binding parameters indicate association (rightward) or dissociation (leftward) **(B)** The simplified model structure from the view of *X*: a feedforward motif, which contains a direct arm from *X* to *EE*, and an indirect arm where *X* acts via the nested HPE feedback loop to affect *EE*.

For simplicity, the hypothalamus and pituitary are lumped into one central module which produces *PH* as the output. The negative feedback action exerted by *EH* on *PH* is modeled as follows. *EH* binds reversibly to the central receptor (*CR*) to form a complex *EHCR*, with a second-order association rate constant *k*_7*f*_ and first-order dissociation rate constant *k*_7*b*_. *PH* is produced at a basal zero-order rate *k*_30_ and an *EHCR*-regulated rate described by an inhibitory Hill function with affinity constant *K*_*d*3_, Hill coefficient *n*_3_, and a maximal synthesis rate constant *k*_3_. The Hill function here provides the ultrasensitivity (percentage-wise amplification) necessary for robust *EH* homeostasis through the negative feedback. *PH* released is degraded in a first-order manner with a rate constant *k*_4_.

To evaluate whether the constructed HPE feedback model can exhibit the typical behaviors of an endocrine feedback system in response to specific perturbations, we first simulated the dynamic responses of *EH* and *PH* in the absence of EDCs. By varying *k*_1_, the rate constant of *PH*-stimulated *EH* production, the model recapitulates the clinical primary hyper- and hypo-functioning endocrine conditions (e.g., primary hyper- or hypothyroidism), where the steady-state *EH* and *PH* levels move in opposite directions, with the fold change of the *PH* level much greater than that of *EH* (Fig. S1A and S1B). By varying *k*_3_, the rate constant of *PH* production, the model recapitulates the clinical secondary hyper- or hypo-functioning conditions, where the steady-state *EH* and *PH* levels move in the same directions with comparable fold changes (Fig. S1C and S1D).

Acting as an agonist or antagonist, an EDC *X* can alter *EE* by competing with *EH* for both the peripheral and central hormone receptors. In the model, *X* can bind reversibly to *PR* in the peripheral module to form a complex *XPR*, with a second-order association rate constant *k*_6*f*_ and first-order dissociation rate constant *k*_6*b*_. When *X* is an agonist, *XPR* is an active complex exerting *EE* with an efficacy *w*_*p*_ relative to *EHPR* (whose efficacy is set at unity) such that the overall *EE* = *EHPR*+*w*_*p*_\**XPR*. When *X* is an antagonist, *XPR* is an inactive complex exerting no *EE*, i.e., *w*_*p*_=0. In the central module, *X* can bind reversibly to *CR* to form a complex *XCR*, with a second-order association rate constant *k*_8*f*_ and first-order dissociation rate constant *k*_8*b*_. When *X* is an agonist, *XCR* is an active complex inhibiting *PH* production with an efficacy *w*_*c*_ relative to *EHCR* (whose efficacy is set at unity) such that the overall inhibitory signaling strength is *EHCR*+*w*_*c*_\**XCR*. When *X* is an antagonist, *XCR* is an inactive complex with *w*_*c*_=0, exerting no inhibition on *PH* production. From the view of *X*, the model forms a feedforward structure, which contains a direct arm from *X* to *EE*, and an indirect arm where *X* acts via the nested HPE feedback loop to affect *EE* (Fig. 1B).

### 2. Construction of a population model of HPE feedback loop

#### 2.1. Normalization of NHANES thyroid profile data

To understand the NMDR behaviors of an ensemble of individuals, we simulated a virtual human population based on the thyroid hormone profiles from the National Health and Nutrition Examination Survey (NHANES) in the three 2-year cycles between 2007-2012. Here the correlated distributions of serum free T4 (fT4) and TSH levels were used to represent the *EH* and *PH* values respectively in the model. After resampling based on normalized sample weight and exclusion of individuals who had taken thyroid drugs, had thyroid cancers, or had missing fT4 or TSH values, the final weight-adjusted population contains 1883 records (details of the process are provide in Supplemental Material).

It was noted that all fT4 levels in cycle I (years 2007-2008) and 57.7% fT4 levels in cycle II (years 2009-2010) of the NHANES dataset were reported with a precision of 0.1 (ng/dL), while the remaining fT4 data including the entire cycle III (years 2011-2012) have a precision of 0.01. To make all fT4 data have a similar fine precision, fT4 levels with a precision of 0.01 in cycles II and III were first merged. fT4 levels in the combined dataset were divided into continuous intervals of 0.05 (ng/dL), and the proportions of samples falling into each interval were calculated. fT4 levels with a precision of 0.1 (ng/dL) in cycles I and II data were imputed based on the calculated sample interval proportions to achieve a precision of 0.01 and the original values were replaced with the imputed values. The final joint distribution of fT4 and Log_10_(TSH) levels were divided into a 28 × 35 grid. fT4 levels were then rescaled to a mean of 10 to represent *EH*, and TSH levels were rescaled to a geometric mean of 1 to represent *PH*. The obtained Log_10_(*PH*) vs *EH* density is shown in Fig. S2.

#### 2.2 Construction of the virtual population HPE model

After normalization with the total count, the density probability of each rectangular unit was either zero or ranged between 0.000052-0.0276. We aimed to produce a virtual population of near 10K individuals. The number of individuals in each unit was determined by multiplying the probability with 10K and rounding to the nearest integer. As a result, a non-empty unit contains a minimum of 5 individuals and maximum of 276 individuals, and the total number of individuals is 9996. To obtain the 10K virtual population model, the minimal HPE model was simulated to steady state by randomly sampling parameter values from log_10_-converted uniform distributions that range between 1/10-10 fold of the default values for *k*_30_, *K*_*d*3_, *k*_7*f*_, and *CR*_*tot*_, between 1/1000-1000 fold of the default value for *k*_1_, and between 1/100-100 fold of the default value for *k*_3_. In addition, *n*_3_ was randomly sampled from a uniform distribution ranging between 4-10. *k*_7*b*_, *k*_2_ and *k*_4_ were not varied because we were only concerned with the steady-state response and varying *k*_7*f*_, *k*_1_ and *k*_3_ was sufficient to achieve this goal. *k*_5*f*_, *k*_5*b*_, and *PR*_*tot*_ were excluded because they are external to the HPE loop. Parameters for *X* binding to *PR* and *CR* were also excluded because they are not part of the HPE axis under physiological conditions.

For each set of randomly sampled parameter values, the HPE model was run and the resulting pair of the steady-state *PH* and *EH* levels was evaluated against the above 28 × 35 grid to determine which rectangular unit the paired values fall into. If the pair fell into a zero-probability unit, the parameter set was rejected. Otherwise, it counts toward the total number of individuals assigned to the unit as above and the corresponding set of parameter values was recorded. When the total number of individuals in a unit was reached, any new pair values of *EH* and *PH* falling in the same unit will be discarded and no parameter values recorded. The random parameter sampling and simulation process continued until all non-zero grid units were filled.

### 3. Classification of NMDR curves

All DR curves were obtained by varying *X* to different values and running the model to steady state. To identify NMDR curves generated by the population model and group similar ones together, the following classification algorithm was used to determine the number of ascending and descending phases in a DR curve. We first calculated the first derivative of each DR curve of *EE*. If the first derivative does not change sign, the DR curve is monotonic; if it changes sign only once, the DR curve is biphasic; if it changes sign twice or more, the DR curve is multiphasic.

### 4. Simulation language and model sharing

Ordinary differential equations (ODEs) describing the rates of change of the state variables are provided in Table S1, and the default parameter values and justifications are provided in Table S2. All simulations and analyses were conducted in MATLAB R2023b (The Mathworks, Natick, Massachusetts, USA). Models were run using *ode23tb* solver to steady state to obtain DR curves unless otherwise indicated. All MATLAB code is available at https://github.com/pulsatility/2024-NMDR-HPE-Model.git.

## Results

### 1. NMDR effects of an EDC agonist

We first explored the situation when an EDC is an agonist, where its molecular action and effect are expected to be in the same direction as the endogenous hormone. In the framework of the HPE model here, a hypothetical EDC agonist, designated as *X* here, acts at both the peripheral and central sites. At the peripheral site, *X* binds to *PR* to form *XPR*, adding to the endocrine effect (*EE*). At the central site, *X* binds to *CR* to form *XCR*, inhibiting *PH* production as would the endogenous *EH* do. The resulting decrease in the *PH* level leads to reduced stimulation of *EH* production and consequently a decrease in the *EH* level and *EHPR*-mediated *EE*. Therefore, the net endocrine outcome of exposure to an agonist *X* depends on the summation of the *XPR*-mediated and *EHPR*-mediated effects, which change in opposite directions as *X* increases.

#### 1.1 Monotonic DR of reference agonist

To establish a reference situation, we first considered when all parameters are at default values and the hypothetical agonist *X* has the same binding affinities and efficacies as *EH* for the two receptors: for *PR*, the dissociation constants *K*_*d*6_ = *K*_*d*5_ (where *K*_*d*6_ = *k*_*b*6_/*k*_*f*6_, *K*_*d*5_ = *k*_*b*5_/*k*_*f*5_) and efficacy *ω*_*p*_ = 1, and for *CR, K*_*d*8_ = *K*_*d*7_ (where *K*_*d*8_ = *k*_*b*8_/*k*_*f*8_, *K*_*d*7_ = *k*_*b*7_/*k*_*f*7_) and efficacy *ω*_*c*_ =1. In this reference situation, *X* is essentially identical to *EH*. As shown in Fig. 2A, as the concentration of *X* increases, more *XCR* is formed. The steady-state *XCR* vs. *X* curve has a Hill coefficient of nearly unity (1.001) and AC_50_ of almost 90 (arbitrary unit, au), which is the same value as the dissociation constant *K*_*d*8_ for the reversible *X* and *CR* binding. Therefore, the *XCR* response is consistent with a receptor-mediated process exhibiting typical Michaelis-Menten kinetics. Increasing *XCR* results in more inhibition of *PH* production and thus a decrease in the steady-state *PH* level (Fig. 2B). When *X* concentration is near 20, *PH* decreases to a basal level. Since *PH* stimulates the production of *EH*, the steady-state *EH* level follows a similar downtrend (Fig. 2C). As a result of the declining *EH*, the steady-state *EHCR* level also decreases (Fig. 2D).

**Figure 2.**
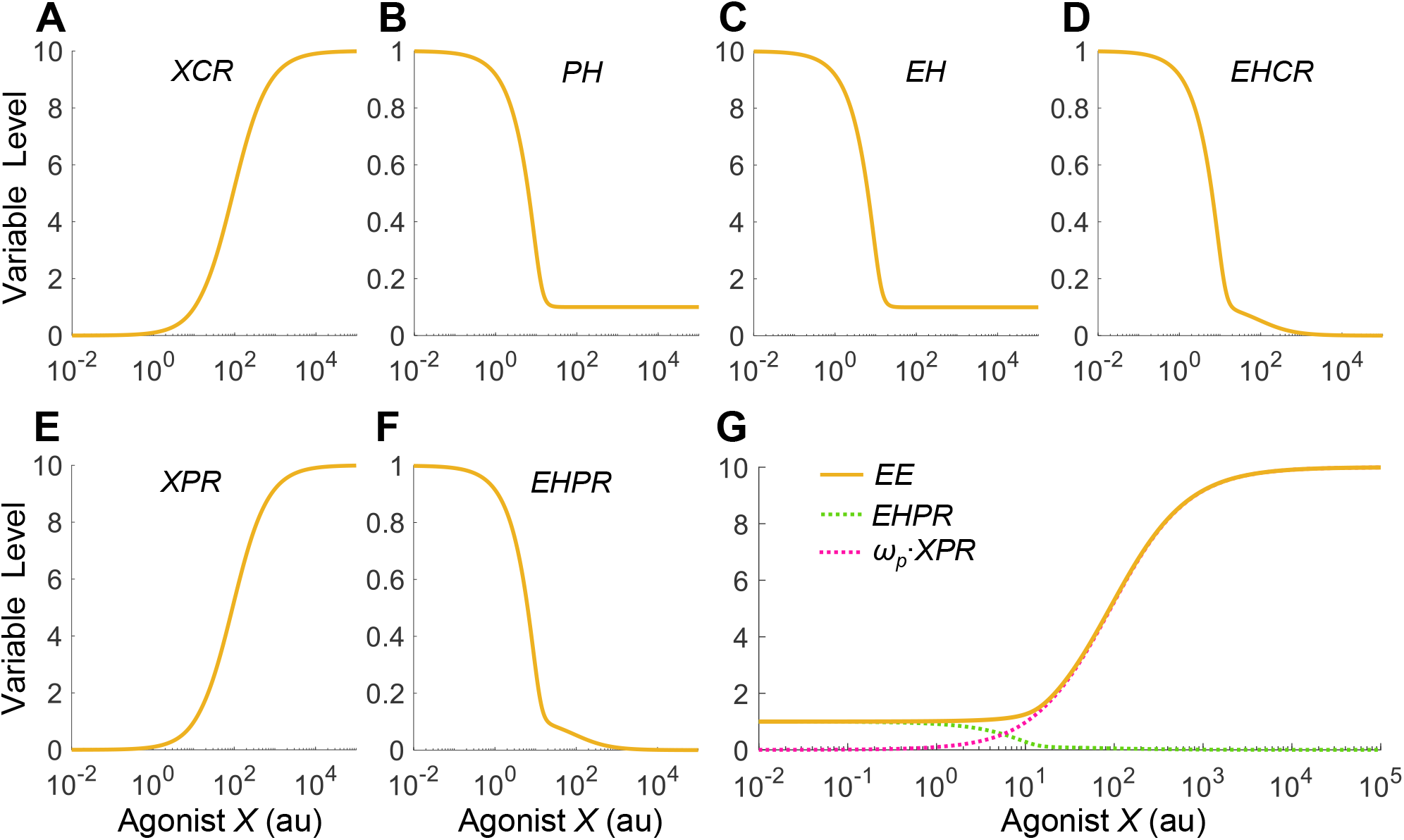
Steady-state DR profiles when *X* acts as a reference agonist. *X* has identical binding affinities and efficacies as *EH* for *CR* and *PR*. Variable names are as indicated. In this reference scenario, the *EE* vs. *X* DR relationship only increases monotonically.

However, *EHCR* also exhibits a secondary decline at *X* concentrations higher than 20 despite that *EH* no longer decreases. This secondary decline occurs because *X* at higher concentrations starts to displace *EH* out of *EHCR* appreciably to form more *XCR* (Fig. 2A). In a similar manner, the declining *EH* also results in a downtrend of the steady-state *EHPR* level with a secondary decline (Fig. 2F), due to competition from the still increasing formation of *XPR* at higher *X* concentrations (Fig. 2E). Lastly, the steady-state *EE* level, which is determined by *EHPR* + *ω*_*p*_\**XPR* (where *ω*_*p*_=1 at default here), exhibits a monotonically increasing, saturable DR profile with respect to *X* (Fig. 2G). The *X*_0.9_/*X*_0.1_ ratio, a metric of the steepness of the curve, is about 45, corresponding to a Hill coefficient of 1.15, which is slightly steeper than the typical receptor-mediated Michaelis-Menten kinetics as exemplified by *XCR* (Fig. 2A) and *XPR* (Fig. 2E).

To further analyze and understand the shape of the *EE* DR curve, we conducted an in-depth analysis of the HPE negative feedback loop. A well-known property of negative feedback is that if the feedback regulation is integral or proportional with high loop gain (amplification), the input-output relationship can be linearized (Zhang and Andersen 2007, Nevozhay, Adams et al. 2009, Sturm, Orton et al. 2010). Considering *X* acting through *CR* as the input and *EH* as an output of the HPE feedback loop, the steady-state *EH* vs. *X* DR relationship indeed follows a nearly straight line that decreases to the basal level on dual-linear scale (Fig. S3B), when the Hill coefficient *n*_3_ (representing the degree of signal simplification, or ultrasensitivity, of the *CR*-mediated feedback here) assumes very high values (*n*_3_ = 1000 or 100). For the high *n*_3_ cases, every increment of *X* concentration results in an almost equal decrement of *EH* concentration, such that the *X + EH* sum remains constant. This occurs because *X*, as the reference agonist here, is parameterized to be indistinguishable from *EH* with identical receptor binding affinity and efficacy properties, such that near-perfect adaption occurs for *X + EH* as a whole. This leads to a flat *EE* response for low *X* levels (Fig. S3D), and an overall monotonically increasing response that is slightly steeper than would be predicted by Michaelis-Menten kinetics (Fig. S3C). Compared with the high *n*_3_ values that can achieve nearly perfect linear *EH* response, substantially lower *n*_3_ values, including the default value of 7, can only achieve partial linearization (Fig. S3B). As *X* increases, *EH* does not decrease as much to match the increase of *X* before bottoming at the basal level. As a result, *XPR* rises faster (Fig. S4E) than *EHPR* declines (Fig. S4F), and *EE*, which equals *EHPR* + *ω*_*p*_·*XPR* (where *ω*_*p*_=1 for the reference situation), can only monotonically increase (Fig. S4G and S3D). In summary, for an agonist that is essentially identical to the endogenous hormone in receptor binding and downstream signaling properties, no nonmonotonic endocrine effect is expected to arise out of the HPE feedback operation, even when the feedback-mediated adaptation is perfect.

#### 1.2 J-shaped DR of agonist – effects of binding affinities for *PR* (*K*_*d*5_ and *K*_*d*6_)

With the reference response established above, we next explored situations when the agonist *X* is quantitatively different than *EH* in receptor binding and efficacy. We first examined the effect of the binding affinity between *X* and *PR* by varying the association rate constant *k*_*f*6_. Since this binding event is outside the HPE feedback loop, the effects of *X* on the components within the feedback loop, including *CR, XCR, EHCR, PH* and *EH*, are the same as the reference situation as in Fig. 2A-2D (results not shown). When the binding affinity between *X* and *PR* is lowered by decreasing *k*_*f*6_, the *XPR* vs. *X* curve shifts to the right as expected, and conversely when *k*_*f*6_ is increased the curve shifts to the left (Fig. 3A). This shift only affects the second decline phase of the *EHPR* response (Fig. 3B), while the first phase remains largely unchanged as it is determined mainly by the declining *EH*. Interestingly, when *k*_*f*6_ is decreased such that *K*_*d*6_ > *K*_*d*5_ appreciably, a J-shaped DR relationship begins to emerge for *EE* (Fig. 3C). At 1/4 of the default value of *k*_*f*6_, *EE* can dip to nearly 50% of the basal level for *X* concentration between 10-20 au (Fig. 3C inset). In contrast, increasing *k*_*f*6_ does not result in a nonmonotonic response. Increasing *K*_*d*6_ by increasing the dissociation rate constant *k*_*b*6_ achieves a similar NMDR effect (results not shown).

**Figure 3.**
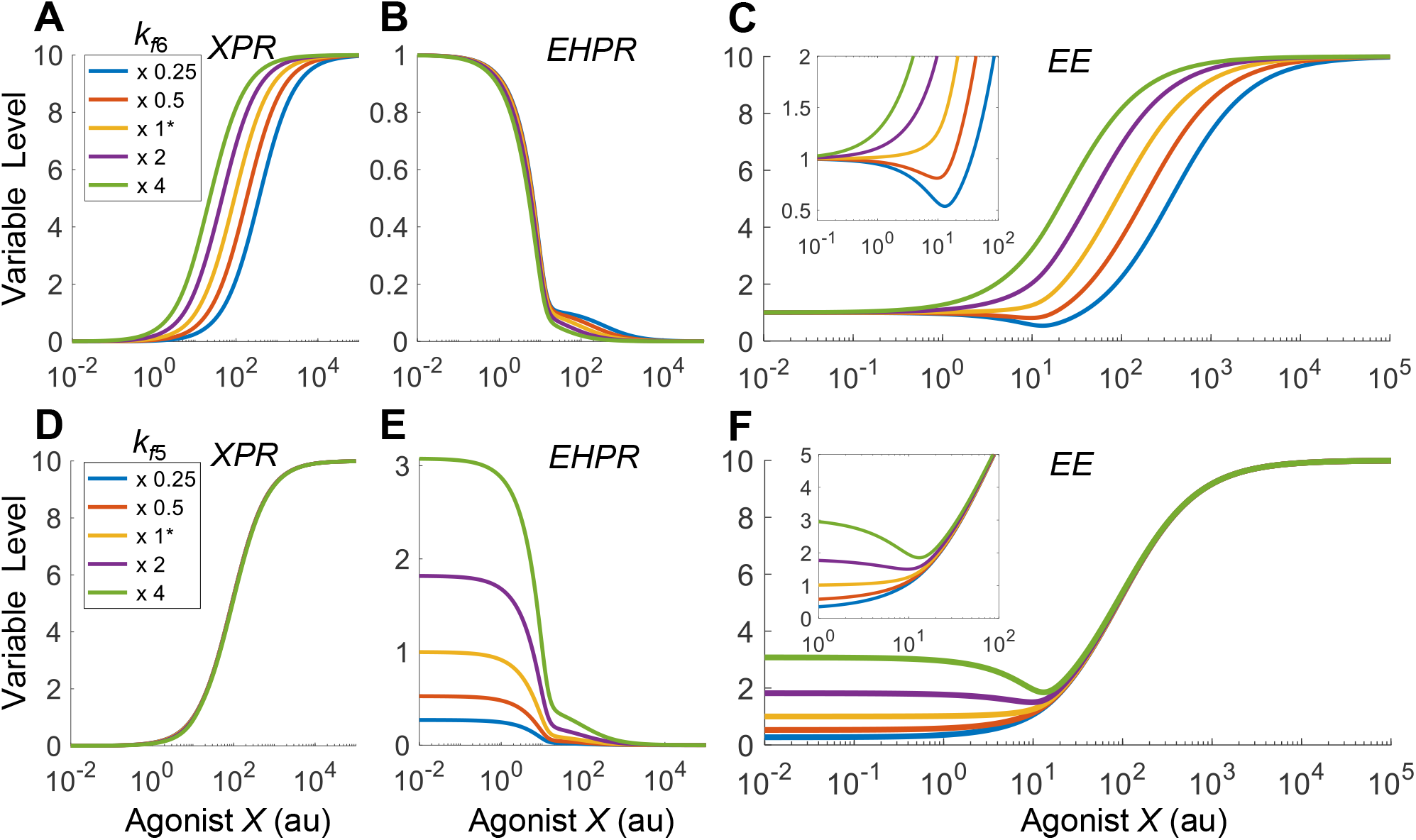
The emergence of J-shaped DR of *EE* when the relative binding affinities of agonist *X* and *EH* for *PR* are different. **(A-C)** J-shaped *EE* response emerges when the binding affinity between *X* and *PR* is decreased by decreasing *k*_*f*6_ from the default value as indicated. **(D-F)** J-shaped *EE* response emerges when the binding affinity between *EH* and *PR* is increased by increasing *k*_*f*5_ from the default value as indicated. Insets: zoomed-in views of the DR curves of *EE* in (C) or (F). x 1* denotes that the parameter is at default value, and x 0.25, x 0.5, x 2, and x 4 denote that the parameter is set at the corresponding fold of the default value. Same denotation is used in other figures where applicable.

The NMDR effect is due to the shift of the *XPR* response curve, which alters the relative contributions of *XPR* and *EHPR* to *EE*. Here the contributions to the change of *EE* are determined by *d(ω*_*p*_\**XPR)/dX* and *dEHPR/dX*, i.e., the slopes of the *ω*_*p*_\**XPR* and *EHPR* curves, respectively (Fig. S5A). As *k*_*f*6_ decreases such that *XPR* gradually shifts to the right, its contribution to the change of *EE* becomes less while the contribution by *EHPR* becomes more dominant. Therefore, for low *k*_*f*6_ values, the *EE* curve initially follows the downtrend of the *EHPR* response at low *X* concentrations. As *X* continues to increase, the slope of the *XPR* curve starts to contribute more than *EHPR* to the change of *EE*, thus the downtrend of the *EE* curve ceases and in turn it starts to rise following the uptrend of the *XPR* response (Fig. S5A). The lower the *k*_*f*6_ value, the higher the magnitude (defined as the vertical drop from the basal *EE* level to the nadir) of the J-shaped effect. The *X* concentration corresponding to the nadir also shifts slightly more to the right but stays in the vicinity of 10-20 au, corresponding to the *X* level when *EH* ceases to decline.

We next examined the effect of the binding affinity between *EH* and *PR* by varying the association rate constant *k*_*f*5_. Since this binding event is also outside the HPE feedback loop, the effects of *X* on the components within the feedback loop are the same as the reference situation as in Fig. 2A-2D (results not shown). When the binding affinity between *EH* and *PR* is increased by increasing *k*_*f*5_, the basal level of *EHPR* is elevated as expected, and the low-dose region of the *EHPR* vs. *X* curve expands upward, and conversely when *k*_*f*5_ is decreased the opposite occurs to the *EHPR* curve (Fig. 3E), without affecting the *XPR* response (Fig. 3D). Interestingly, when *k*_*f*5_ is increased such that *K*_*d*6_ > *K*_*d*5_ appreciably, a J-shaped DR relationship begins to emerge for *EE* (Fig. 3F). In contrast, decreasing *k*_*f*5_ does not result in a nonmonotonic response. Decreasing *K*_*d*5_ by decreasing the dissociation rate constant *k*_*b*5_ achieves a similar NMDR effect (results not shown). Just as the case of varying *K*_*d*6_, varying *K*_*d*5_ changes the relative contribution of *EHPR* and *XPR* to *EE* (Fig. S5B). J-shaped DR emerges when the binding affinity between *EH* and *PR* is high (i.e., low *K*_*d*5_), where the contribution by *EHPR* to the change of *EE* is greater than the contribution by *XPR* at low *X* concentrations.

#### 1.3 J-shaped DR of agonist – effects of binding affinities for *CR* (*K*_*d*7_ and *K*_*d*8_)

We first examined the effect of the binding affinity between *X* and *CR*. When the binding affinity is lowered by decreasing the association rate constant *k*_*f*8_, the *XCR* vs. *X* curve shifts to the right as expected, and conversely when *k*_*f*8_ is increased the curve shifts to the left (Fig. 4A). Through the inhibitory action of *XCR* on *PH*, this shift propagates downstream, leading to similar shifts of the *PH, EH, EHCR*, and *EHPR* responses (Fig. 4B-4D, 4F), without affecting the *XPR* response (Fig. 4E). When *k*_*f*8_ is increased such that *K*_*d*7_ > *K*_*d*8_ appreciably, J-shaped DR relationships begin to emerge for *EE* (Fig. 4G). Decreasing *K*_*d*8_ by decreasing the dissociation rate constant *k*_*b*8_ achieves a similar NMDR effect (results not shown). The horizontal shift of the *EHPR* response changes its relative contribution to *EE* (Fig. S5C), and a J-shaped DR of *EE* emerges when the binding affinity between *X* and *CR* is high (i.e., low *K*_*d*8_), where the contribution to the change of *EE* by *EHPR* dominates that by *XPR* at low *X* concentrations.

**Figure 4.**
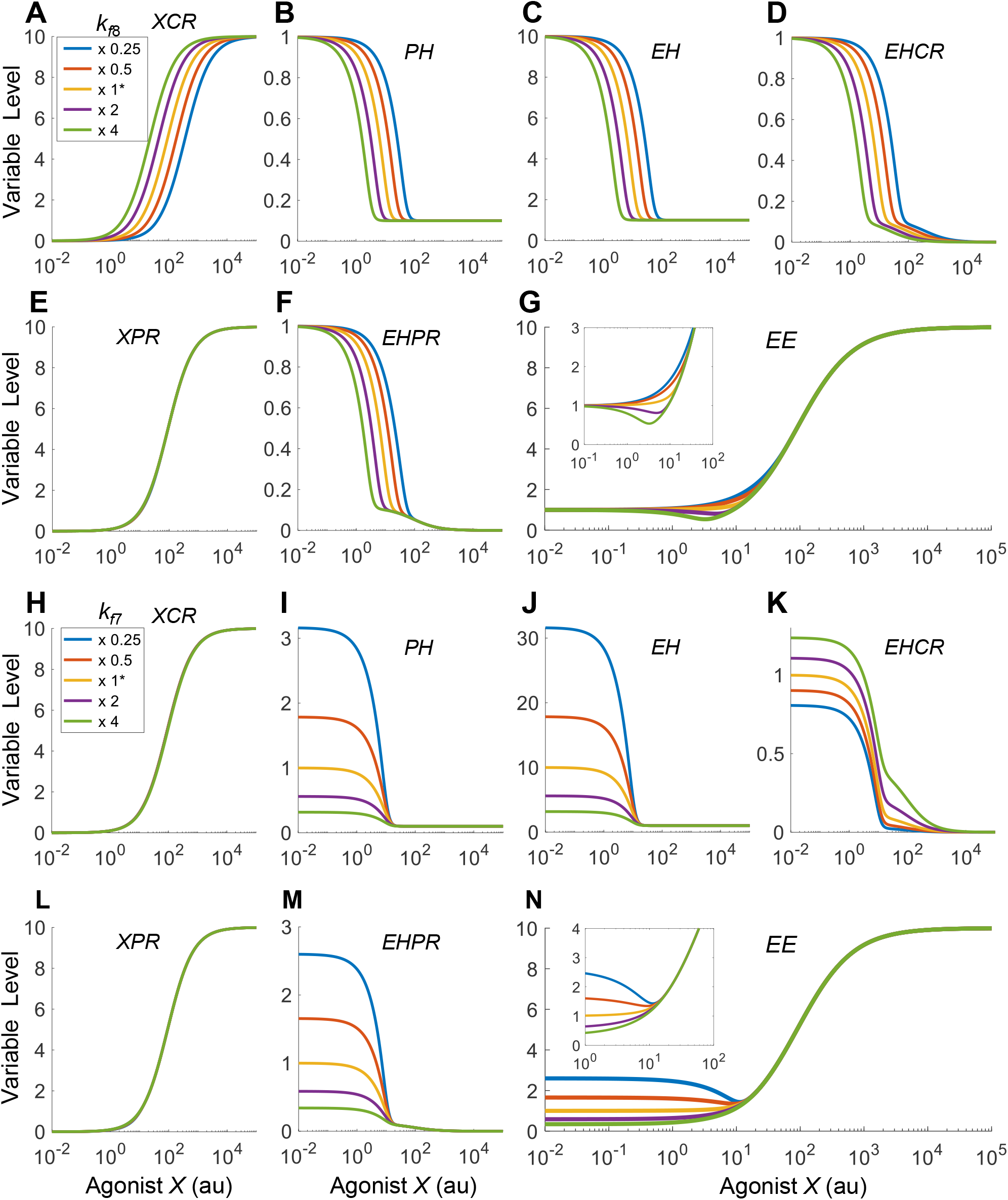
The emergence of J-shaped DR of *EE* when the relative binding affinities of agonist *X* and *EH* for *CR* are different. **(A-G)** J-shaped *EE* response emerges when the binding affinity between *X* and *CR* is increased by increasing *k*_*f*8_ from the default value as indicated. **(H-N)** J-shaped *EE* response emerges when the binding affinity between *EH* and *CR* is decreased by decreasing *k*_*f*7_ from the default value as indicated. Insets: zoomed-in views of the DR of *EE* in (G) or (N).

We next examined the effect of the binding affinity between *EH* and *CR*. When the binding affinity is increased by increasing the associate rate constant *k*_*f*7_, the feedback inhibition by *EH* on *PH* is enhanced, and as a result the basal *PH* level decreases and the opposite occurs when *k*_*f*7_ is decreased (Fig. 4I). This effect propagates downstream to *EH* and *EHPR* (Fig. 4J and 4M). Interestingly, *EHCR* exhibits an opposite effect with much smaller changes in the basal level (Fig. 4K). This is because as *k*_*f*7_ increases, the increased *EHCR* formation is partially cancelled out by the decreasing *EH*. The *XCR* and *XPR* responses are not altered by *k*_*f*7_ (Fig. 4H and 4L). When *k*_*f*7_ is decreased such that *K*_*d*7_ > *K*_*d*8_ appreciably, J-shaped DR relationships begin to emerge for *EE* (Fig. 4N). Increasing *K*_*d*7_ by increasing the dissociation rate constant *k*_*b*7_ achieves a similar NMDR effect (results not shown). Varying *K*_*d*7_ changes the relative contribution of *EHPR* (as its basal level changes dramatically) to *EE* (Fig. S5D). J-shaped DR emerges when the binding affinity between *EH* and *CR* is low (i.e., high *K*_*d*7_), where the contribution to the change of *EE* by *EHPR* dominates that by *XPR* at low *X* concentrations.

In summary, the simulation results above indicate that when the efficacies of *X* and *EH* are comparable (*ω*_*p*_ ≈ *ω*_*c*_ ≈ 1), as long as the relative binding affinity of *X* for *CR* vs. *X* for *PR* (defined as *K*_*d*6_/*K*_*d*8_) is appreciably greater than the relative binding affinity of *EH* for *CR* vs. *EH* for *PR* (defined as *K*_*d*5_/*K*_*d*7_), a J/U-shaped DR relationship for *EE* would emerge.

#### 1.4 J-shaped DR of agonist – effects of efficacy (*ω*_*p*_ and *ω*_*c*_)

We next explored whether the efficacy of *X* also plays a role in determining the nonmonotonic effect. We first examined the *ω*_*p*_, the efficacy of *X* acting via *XPR* to produce *EE*. Since *ω*_*p*_ is a parameter outside of the HPE feedback loop, the effects of *X* on the variables within the feedback loop are the same as the reference situation as in Fig. 2A-2D (results not shown), so are the *XPR* and *EHPR* responses (Fig. 5A and 5B) since the receptor binding *per se* is not affected by *ω*_*p*_. However, because varying *ω*_*p*_ alters the relative contribution of *XPR* to *EE*, a J-shaped DR for *EE* emerges when *ω*_*p*_ is tangibly smaller than unity which is the efficacy of *EH* for *PR* (Fig. 5C inset and Fig. S5E).

**Figure 5.**
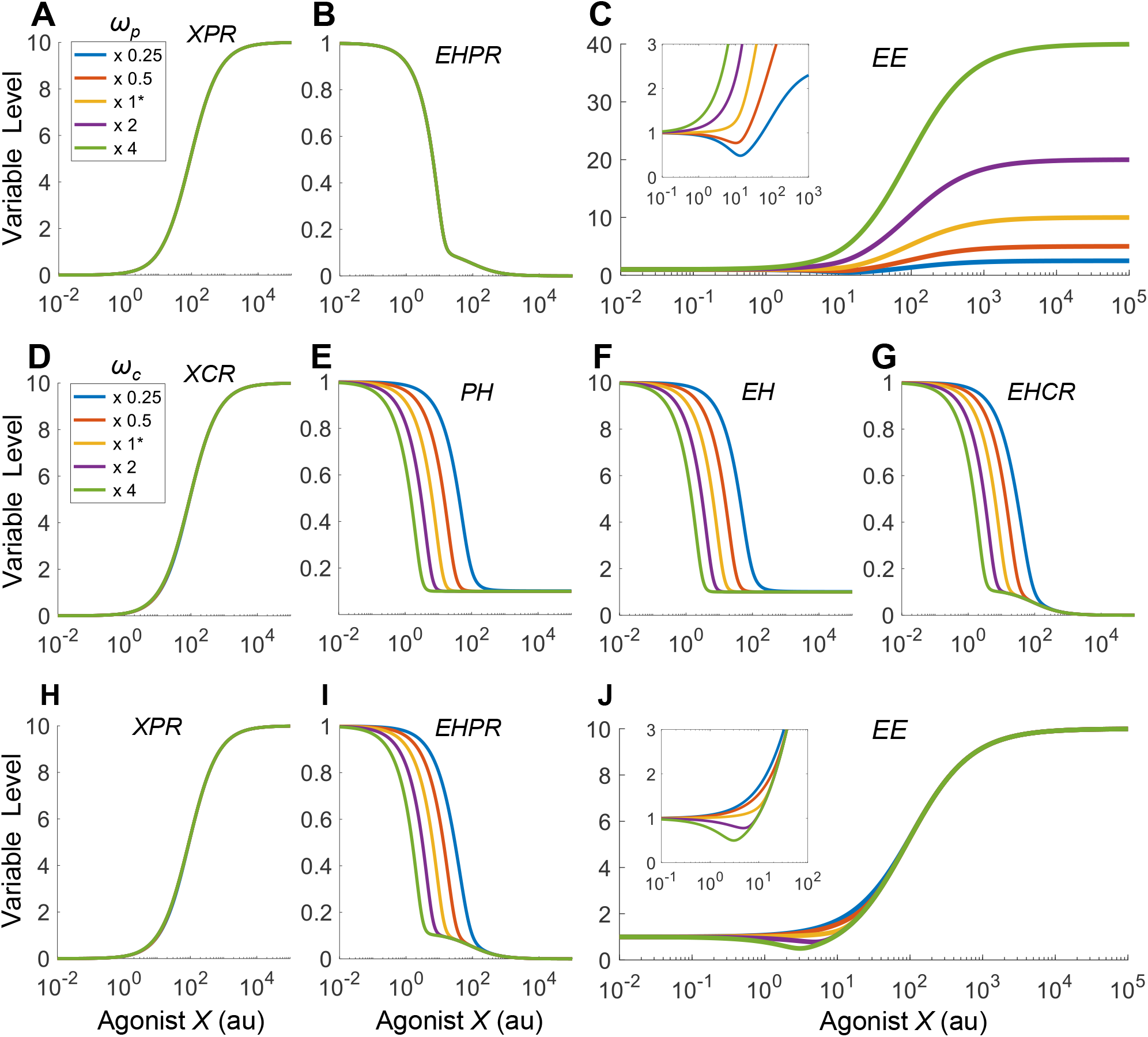
The emergence of J-shaped DR of *EE* when the efficacies of agonist *X* and *EH* are different. **(A-C)** J-shaped *EE* response emerges when the efficacy of *XPR* (*ω*_*p*_) is decreased from the default value as indicated. **(D-J)** J-shaped *EE* response emerges when the efficacy of *XCR* (*ω*_*c*_) is increased from the default value as indicated. Insets: zoomed-in views of the DR of *EE* in (C) or (J).

We next examined *ω*_*c*_, the efficacy of *X* acting via *XCR* to inhibit *PH* production. When *ω*_*c*_ is lowered, although the *XCR* vs. *X* curve is not affected (Fig. 5D), the inhibition of *PH* by *XCR* is reduced, causing the *PH* vs. *X* curve to shift to the right, and conversely when *ω*_*c*_ is increased the curve shifts to the left (Fig. 5E). This shift propagates downstream, leading to similar shifts of the *EH, EHCR*, and *EHPR* responses (Fig. 5F-5G, 5I), without affecting the *XPR* response (Fig. 5H). The horizontal shift of the *EHPR* response changes its relative contribution to *EE*, and a J-shaped DR emerges when *ω*_*c*_ is tangibly greater than unity which is the efficacy of *EH* for *CR* (Fig. 5J inset and Fig. S5F).

#### 1.5 Monotonic DR of agonist - effects of remaining parameters

Lastly, we examined the effects of the remaining parameters, including *k*_1_, *k*_2_, *k*_3_, *k*_4_, *k*_30_, *K*_d3_, *n*_3_, *CR*_*tot*_, and *PR*_*tot*_. We found that even though the value of each of these parameters was varied by 0.01-100 fold, no NMDR emerges (simulation results not shown). Taken together, these results indicate that only the six parameters related to receptor binding affinity and efficacy play a role in rendering NMDR under the current parameter condition.

### 2. NMDR effects of an EDC antagonist

When an EDC is an antagonist, it is still capable of receptor binding, but the binding does not lead to downstream molecular action. In the framework of the HPE model here, antagonist *X* is mimicked by setting the efficacy parameters *ω*_*p*_ and *ω*_*c*_ to zero as the default condition, such that *X* only competes with *EH* for *PR* and *CR* binding, but producing no downstream *XPR*-mediated *EE* and *XCR*-mediated feedback inhibition of *PH*. As a result, at the peripheral site, by sequestering *PR* and displacing *EH* out of the *EHPR* complex, *X* tends to reduce *EE*. At the central site, by displacing *EH* out of the *EHCR* complex, *X* relieves *EHCR*-imposed inhibition of *PH* production, leading to increased *PH* and subsequently increased *EH* levels. Therefore, the net endocrine outcome of exposure to antagonist *X* will depend on the mathematical product of two opposing changes, a declining free *PR* and an increasing *EH*, as the two bind together to form *EHPR* that produces *EE*.

#### 2.1 Monotonic DR of reference antagonist

To establish a reference point for the antagonist, we first considered the baseline situation where all parameters are at default values and the hypothetical antagonist *X* has the same binding affinities for the receptors as *EH*, i.e., *K*_*d*6_ = *K*_*d*5_ and *K*_*d*8_ = *K*_*d*7_, except that the efficacies *ω*_*p*_ = 0 and *ω*_*c*_ = 0. As shown in Fig. 6A, as the concentration of *X* increases, more *XCR* is formed. The steady-state *XCR* vs. *X* curve has a Hill coefficient of 0.88 and AC_50_ of nearly 111, which is slightly higher than *K*_*d*8_ for the *X* and *CR* binding event. Therefore, the *XCR* response is slightly more subsensitive than typical Michaelis-Menten kinetics, and so is the *XPR* response (Fig. 6E). By sequestering *CR* to form *XCR, X* drives *EHCR* to lower levels (Fig. 6D), resulting in less inhibition of *PH* production and thus higher *PH* (Fig. 6B) and *EH* (Fig. 6C) levels. When *X* concentration is near 3000, *PH* and *EH* hit the plateaus at maximum. Similarly, by sequestering *PR* to form *XPR* (Fig. 6E), *X* drives free *PR* to lower levels (Fig. 6G). Despite that *EH* rises as *X* increases, the formation of *EHPR* and thus *EE* continue to decrease monotonically (Fig. 6H). The Hill coefficient of the *EE* vs. *X* DR curve is 0.94.

**Figure 6.**
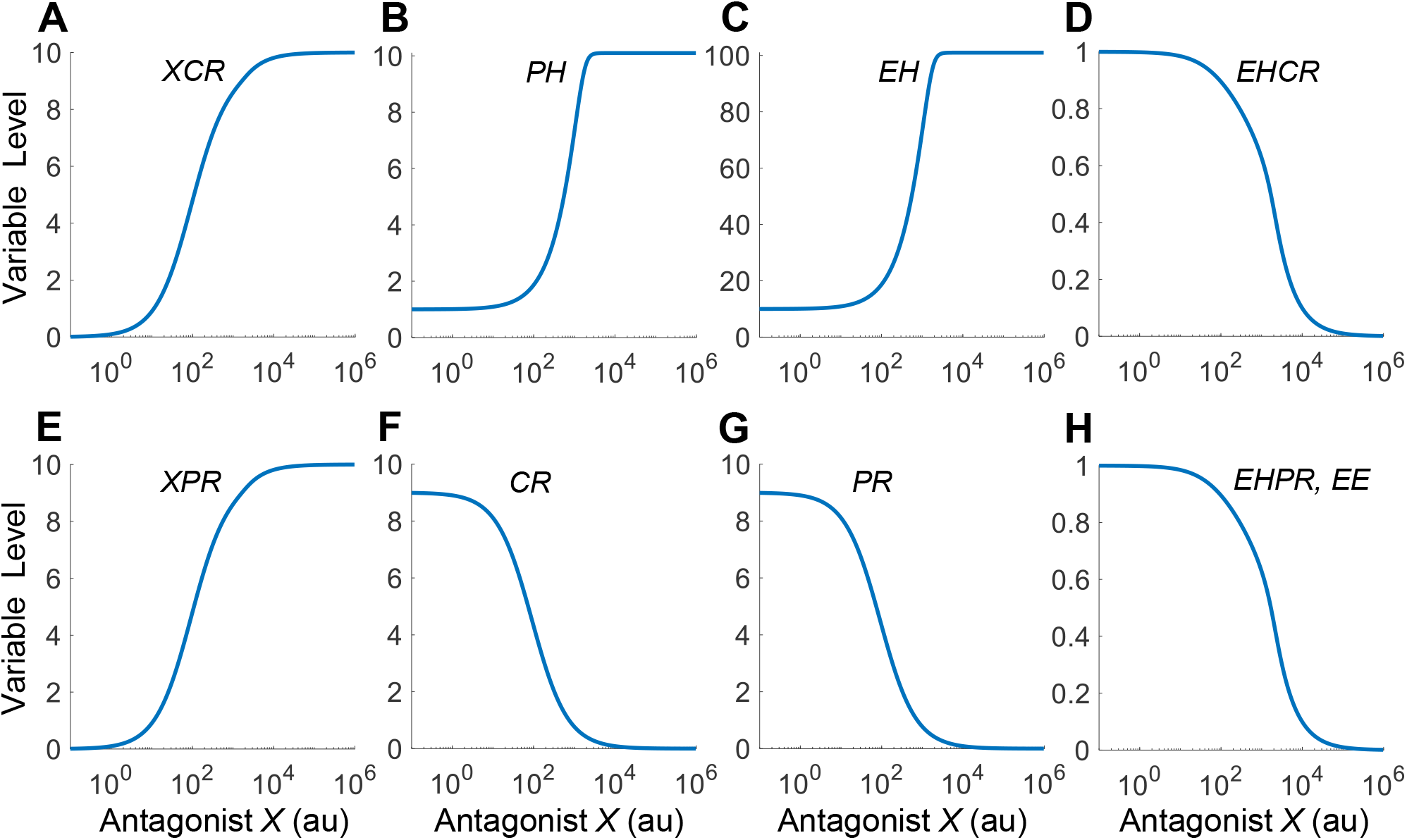
Steady-state DR profiles when *X* acts as a reference antagonist. *X* has identical binding affinities as *EH* for *CR* and *PR* but the efficacies are zero. Variable names are as indicated. In this reference scenario, the *EE* vs. *X* DR relationship only decreases monotonically.

As in the agonist case, we conducted an analysis of the HPE negative feedback loop for antagonist *X* at different degrees of signal simplification. When the Hill coefficient *n*_3_ is at very high values, the steady-state *EH* vs. *X* DR relationship is also a nearly linear response, which increases then plateaus on dual-linear scale (Fig. S6B, *n*_3_ = 100 or 1000). Interestingly, this linear increase in *EH* somehow cancels out the effect of decreasing *PR* levels (Fig. S6D), leading to a flat *EE* response for low *X* concentrations (Fig. S6F), and an overall monotonically decreasing *EE* vs. *X* DR curve (Fig. S6E). Compared with the high *n*_3_ values that can achieve nearly perfect linear *EH* vs. *X* response, substantially lower *n*_3_ values, including the default value of 7, can barely achieve linearization (Fig. S6B). As *X* increases, *EH* does not increase as much to cancel out the effect of the decreasing *PR* before plateauing. As a result, *EHPR* (Fig. 6H) and thus *EE* (Figs. 6H, S6E and S6F) can only monotonically decrease. In summary, for a full antagonist that has the same affinity as the endogenous hormone for receptor binding, no nonmonotonic endocrine effect is expected to arise out of the HPE feedback operation.

#### 2.2 Bell-shaped DR of antagonist – effects of binding affinities for *PR* (*K*_*d*5_ and *K*_*d*6_)

With the reference response established above, we next explored situations when the antagonist *X* is quantitatively different than the endogenous hormone in receptor binding affinity. We first examined the effect of the binding affinity between *X* and *PR* by varying the association rate constant *k*_*f*6_ (Fig. 7A-7C). Since this binding event is outside of the HPE feedback loop, the effects of *X* on the components within the feedback loop, including *CR, XCR, EHCR, PH* and *EH*, are the same as the baseline situation as in Fig. 6A-6D and 6F (results not shown). When the binding affinity between *X* and *PR* is lowered by decreasing *k*_*f*6_, the *XPR* vs. *X* curve shifts to the right, as expected, and conversely when *k*_*f*6_ is increased the curve shifts to the left (Fig. 7A). This shift also leads to a corresponding shift of the *PR* response in the opposite direction (Fig. 7B). Interestingly, when *k*_*f*6_ is decreased such that *K*_*d*6_ > *K*_*d*5_ appreciably, Bell-shaped DR relationships begin to emerge for *EHPR* and *EE* (Fig. 7C). In contrast, increasing *k*_*f*6_ does not lead to nonmonotonic responses. Increasing *K*_*d*6_ by increasing the dissociation rate constant *k*_*b*6_ achieves a similar NMDR effect (results not shown).

**Figure 7.**
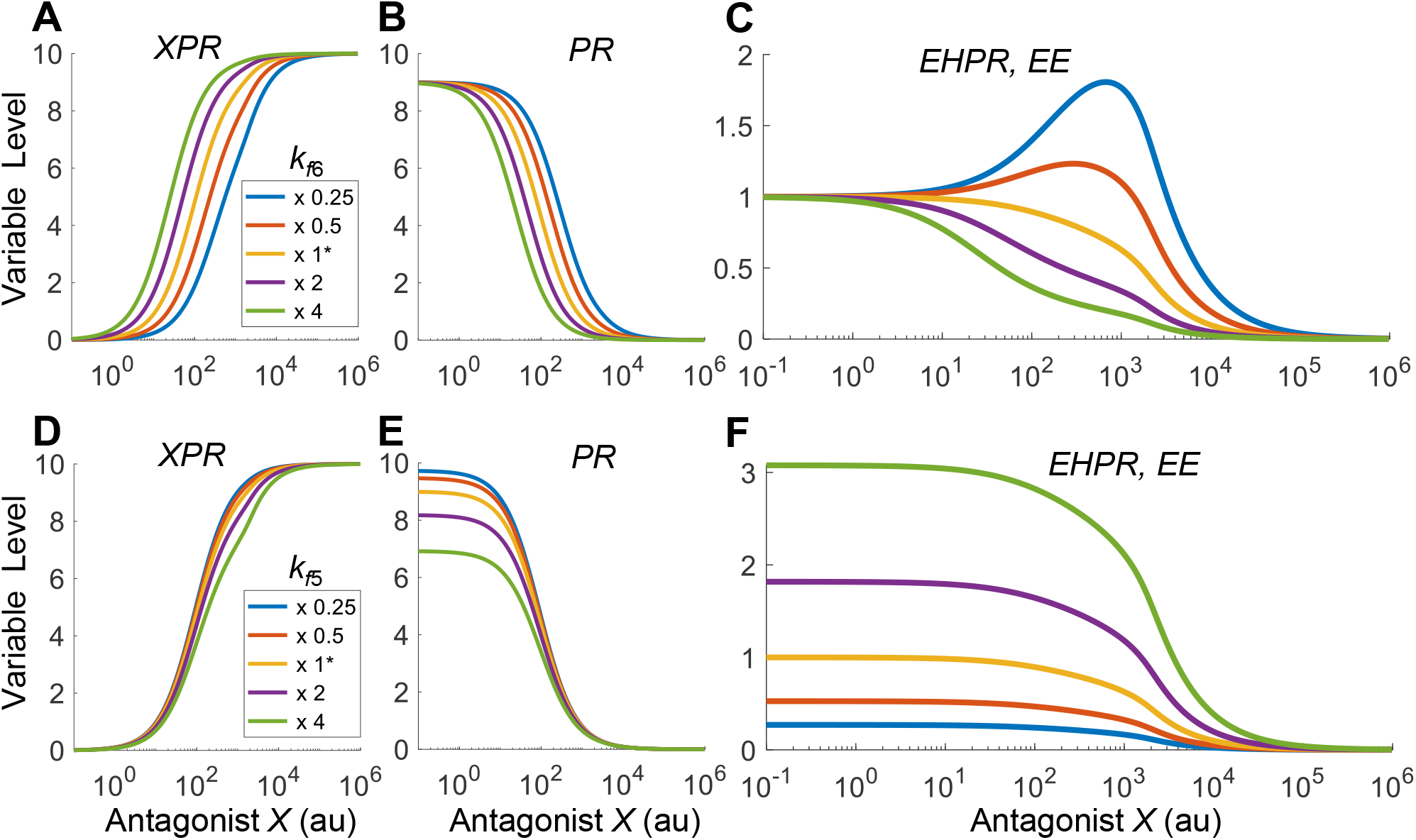
The emergence of Bell-shaped DR of *EE* when the relative binding affinities of antagonist *X* and *EH* for *PR* are different. **(A-C)** Bell-shaped *EE* response emerges when the binding affinity between *X* and *PR* is decreased by decreasing *k*_*f*6_ from the default value as indicated. **(D-F)** Monotonic *EE* responses when the binding affinity between *EH* and *PR* is varied by varying *k*_*f*5_ from the default value as indicated.

The NMDR effect as *K*_*d*6_ is increased is due to the shift of the DR curve of *PR* which alters its relative contributions, compared with *EH*, to *EHPR* formation and thus *EE* response. Here *EHPR* formation is determined by the mathematical product of *PR* and *EH* based on mass action. As *k*_*f*6_ decreases such that *PR* gradually shifts to the right, its contribution to the change in *EHPR* formation, as determined by the slope on the dual-log scale (Fig. S7A), becomes less while the contribution by *EH* becomes relatively more dominant. Therefore, for low *k*_*f*6_ values, the *EE* curve initially follows the uptrend of *EH* at low *X* concentrations. As *X* continues to increase, with *EH* approaching a plateau *PR* starts to influence more, thus the *EE* curve starts to decrease following the downtrend of *PR*. The lower the *k*_*f*6_ value, the higher the magnitude (defined as the vertical elevation from the basal *EE* level to the peak) of the Bell-shaped response, with the peak shifting more to the right.

We next examined the effect of the binding affinity between *EH* and *PR* by varying the association rate constant *k*_*f*5_ (Fig. 7D-7F). Similar to *k*_*f*6_, since this binding event also sits outside of the HPE feedback loop, it does not affect the responses of the components within the loop, which are the same as the baseline situation as in Fig. 6A-6D and 6F (results not shown). When the binding affinity between *EH* and *PR* is increased (decreased) by increasing (decreasing) *k*_*f*5_, the *PR* vs. *X* curve, which monotonically decreases (Fig. 7E) is lowered (elevated) at low *X* concentrations, while the *EHPR* and thus *EE* responses (Fig. 7F) are elevated (lowered). Unlike the agonist case, the *EE* response remains monotonically decreasing as *k*_*f*5_ is varied in either direction. Varying *K*_*d*5_ by varying the dissociation rate constant *k*_*b*5_ does not produce NMDR effects either (results not shown). This lack of NMDR can be traced to the vertical shift of the *PR* vs. *X* curve, without slope changes, as shown on the dual-log scale (Fig. S7B).

#### 2.3 Bell-shaped DR of antagonist – effects of binding affinities for *CR* (*K*_*d*7_ and *K*_*d*8_)

We first examined the effect of the binding affinity between *X* and *CR* (Fig.8A-8H). When the binding affinity is lowered by decreasing the association rate constant *k*_*f*8_, the *XCR* vs. *X* curve shifts to the right, and conversely when *k*_*f*8_ is increased the curve shifts to the left (Fig. 8A). As *X* competes with *EH* for *CR*, this leads to similar shifts of the *EHCR* (Fig. 8D) and *CR* (Fig. 8F) responses. The shift in the *EHCR* response propagates downstream, leading to similar shifts of the *PH* and *EH* responses (Fig. 8B-8C), without tangibly affecting the *XPR* and *PR* responses (Fig. 8E and 8G). When *k*_*f*8_ is increased such that *K*_*d*7_ > *K*_*d*8_ appreciably, Bell-shaped DR relationships begin to emerge for *EHPR* and *EE* (Fig. 8H). Decreasing *K*_*d*8_ by decreasing the dissociation rate constant *k*_*b*8_ achieves a similar NMDR effect (results not shown). The horizontal shift of the *EH* response changes its relative contribution to the formation of *EHPR* and thus *EE* response (Fig. S7C), and Bell-shaped DR emerges when the binding affinity between *X* and *CR* is high (i.e., low *K*_*d*8_), where the contribution to the change in *EE* at low *X* concentrations is dominated by *EH* than by *PR*.

**Figure 8.**
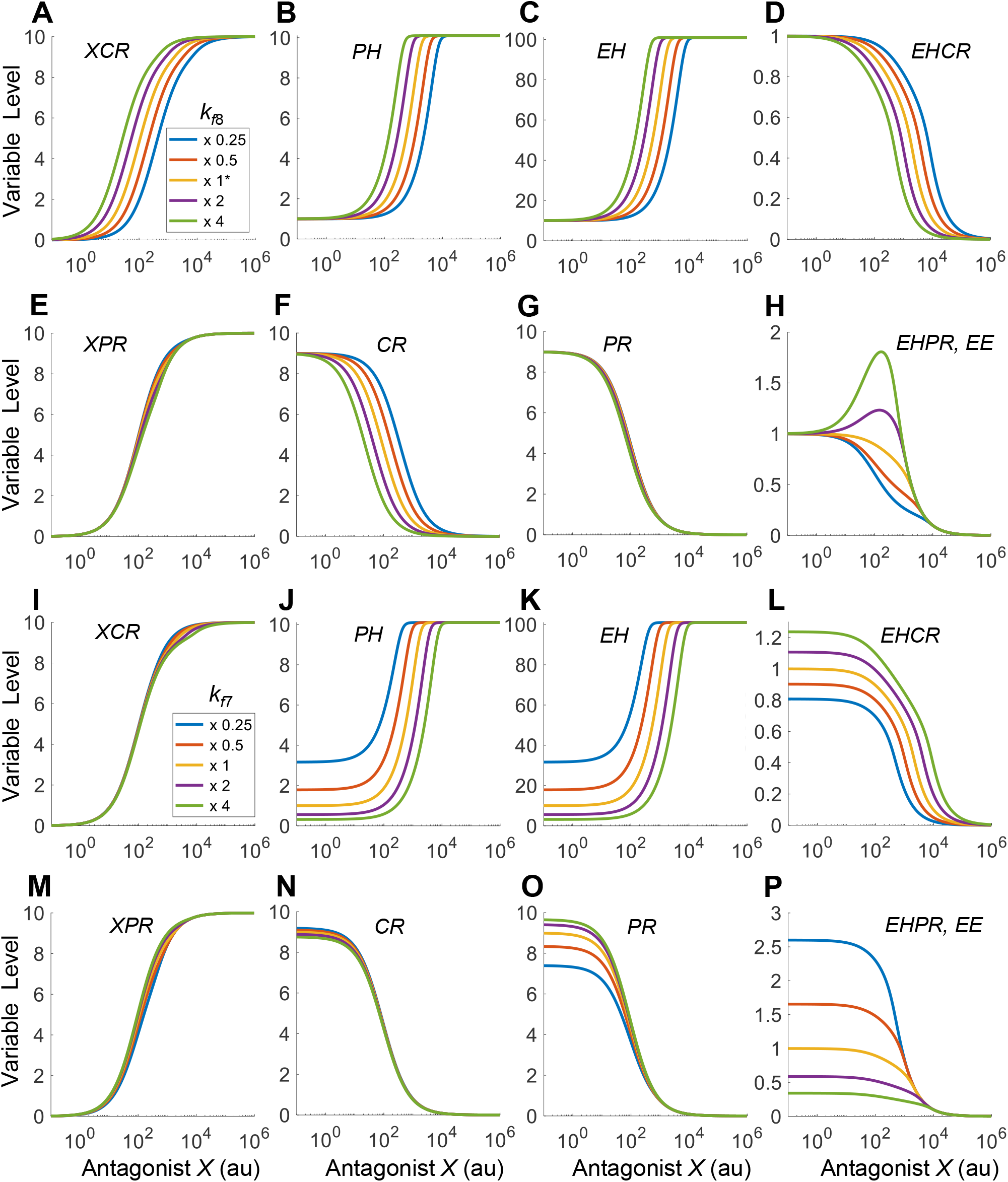
The emergence of Bell-shaped DR of *EE* when the relative binding affinities of antagonist *X* and *EH* for *CR* are different. **(A-H)** Bell-shaped *EE* response emerges when the binding affinity between *X* and *CR* is increased by increasing *k*_*f*8_ from the default value as indicated. **(I-P)** Monotonic *EE* responses when the binding affinity between *EH* and *CR* is varied by varying *k*_*f*7_ from the default value as indicated.

We next examined the effect of the binding affinity between *EH* and *CR* (Fig. 8I-8P). When the binding affinity is increased (decreased) by increasing (decreasing) *k*_*f*7_, the *PH* and *EH* vs. *X* curves, which monotonically increase (Fig. 8J and 8K), is shifted is to the right (left) and the segment at low *X* concentrations is lowered (elevated), while the *EHPR* and thus *EE* responses (Fig. 8P) are elevated (lowered). Unlike the agonist case, the *EE* response remains monotonically decreasing as *k*_*f*7_ is varied in either direction. Varying *K*_*d*7_ by varying the dissociation rate constant *k*_*b*7_ does not produce NMDR effects either (results not shown). This lack of NMDR can be traced to the vertical shift of the segment of the *EH* vs. *X* curve at low *X* concentrations, as shown on the dual-log scale (Fig. S7D).

In summary, the simulations above indicate that only when the binding affinities of *X* for *PR* or *CR* themselves are altered to be different than those of *EH* for *PR* or *CR*, will nonmonotonic endocrine effects emerge. In contrast, varying the binding affinities of *EH* for *PR* or *CR*, which causes changes in the baseline *EE*, does not produce NMDR effects.

#### 2.4 Monotonic DR of antagonist – effects of remaining parameters

Lastly, we examined the effects of the remaining parameters, including *k*_1_, *k*_2_, *k*_3_, *k*_4_, *k*_30_, *K*_d3_, *n*_3_, *CR*_*tot*_, and *PR*_*tot*_. We found that even though the value of each of these parameters was varied by 0.01-100 fold, no NMDR emerges (simulation results not shown).

### 3. NMDR in Monte Carlo simulations

In the above sections, we found that six parameters (*K*_d5_, *K*_d6_, *K*_d7_, *K*_d8_, *ω*_*p*_ and *ω*_*c*_) or their subset play a role in rendering J/U-shaped or Bell-shaped (as opposed to monotonically increasing or decreasing) responses to an agonist or antagonist. In section we further explored the quantitative relationships between these parameters that enable NMDR. We hypothesize that for a J/U-shaped NMDR to occur, the relationship between these six parameters needs to meet the following condition, under the assumption that *X* has the same concentration in the peripheral target tissue as in the brain, and so does *EH* (see Discussion for scenarios of differential concentrations at different site):

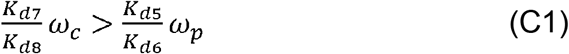

C1 indicates that the central action of *X* relative to *EH* to inhibit *PH* needs to be greater than the peripheral action of *X* relative to *EH* to produce *EE*.

For a Bell-shaped NMDR to occur, the relationship between the six parameters needs to meet the following condition, also under the assumption that *X* has the same concentration in the peripheral target tissue as in the brain, and so does *EH*:

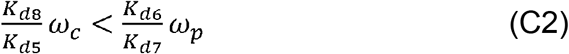

C2 indicates that the central action of *X* to block *EH*-mediated negative feedback thus disinhibiting *PH* needs to be greater than the peripheral action of *X* to block *EH*-mediated *EE*.

To validate these conditions in a more unbiased manner, we conducted Monte Carlo (MC) simulations with two different approaches: (i) randomizing the values of these 6 parameters only while holding other parameters at default values, (ii) utilizing a population HPE model where all relevant model parameters are different between individuals.

#### 3.1. Six-Parameter MC simulations

20,000 MC simulations were conducted to generate a variety of shapes of DR curves by simultaneously sampling (i) parameters *K*_d5_, *K*_d6_, *K*_d7_, and *K*_d8_ from log_10_ uniform distributions ranging between 10-fold above and below default values, and (ii) parameters *ω*_*p*_ and *ω*_*c*_ from uniform distributions of [0-1]. After classification, the DR curves fall into 4 categories: monotonically increasing (MI), J/U, monotonically decreasing (MD), and Bell shapes (Fig. S8A-S8D). The percentage distributions of these different shapes are 49, 45, 3, and 3% respectively (Fig. S8E). Therefore, MI and J/U curves are ~16 times more frequent than MD and Bell curves when only the 6 parameters are randomly sampled.

Among the MI and J/U curves, the paired values of 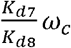 and 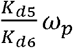 are distributed predominantly above the diagonal for J/U curves, and below the diagonal for MI curves (Fig. 9A).

**Figure 9.**
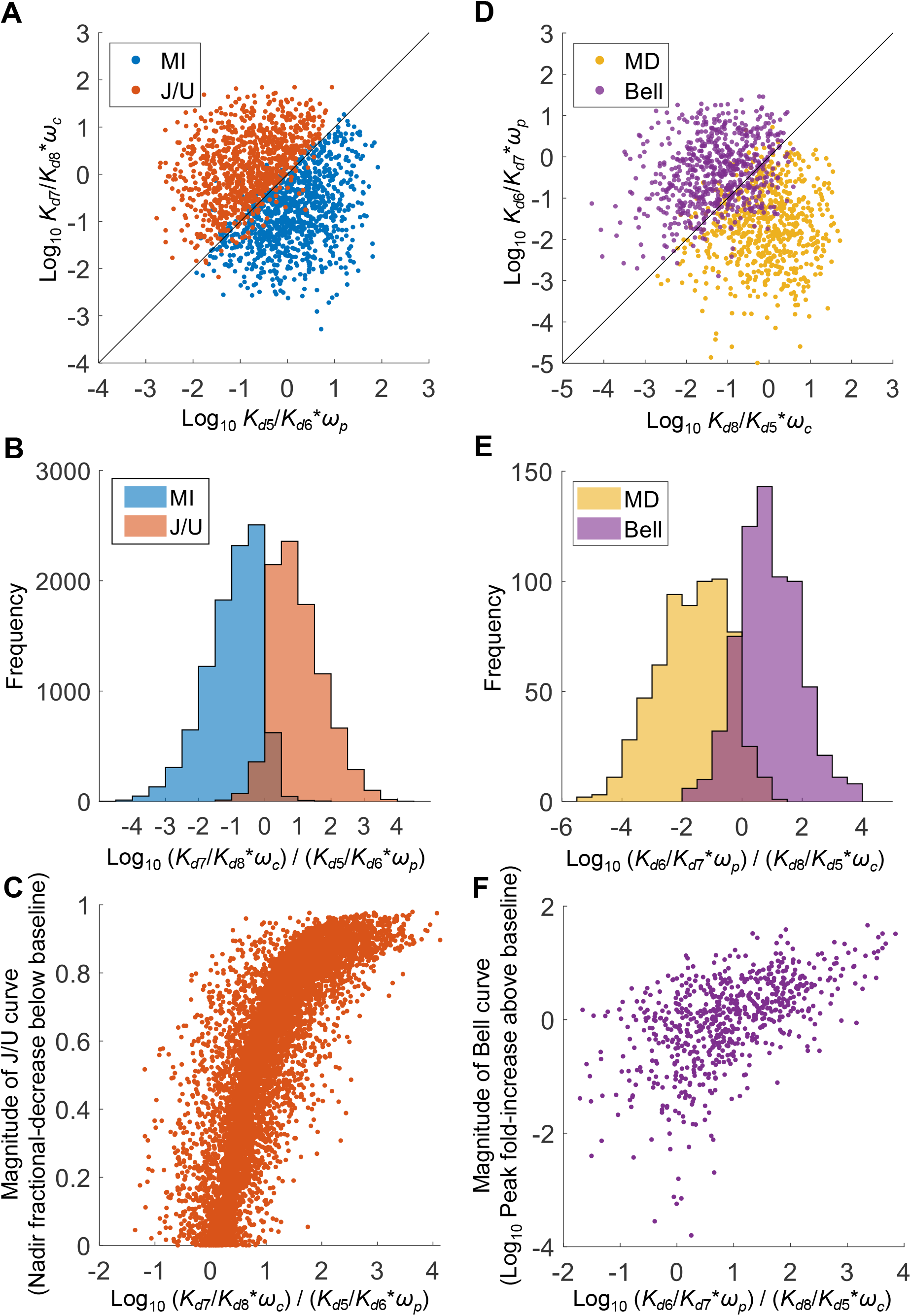
Relationships between parameters *K*_*d*5_, *K*_*d*6_, *K*_*d*7_, *K*_*d*8_, *ω*_*p*_, and *ω*_*c*_ for MI vs. J/U and MD vs. Bell curves from 20,000 six-parameter MC simulations. *k*_5_, *k*_6_, *k*_7_, and *k*_8_ were randomly sampled from uniform distributions of log_10_([0.1, 10]) as fold change relative to the respective default values, and *ω*_*p*_ and *ω*_*c*_ were randomly sampled from the uniform distribution [0,1]. For MI and J/U curves, **(A)** scatter plot of logarithmic 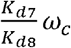 vs. 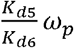 of 1000 randomly selected paired values, **(B)** distribution histograms of logarithmic 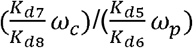, **(C)** scatter plot of magnitude of J/U curves, defined as the fractional-decrease of nadir *EE* from the baseline level when *X*=0, vs. logarithmic 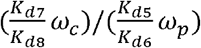. For MD and Bell curves, **(D)** scatter plot of logarithmic 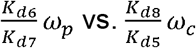, **(E)** distribution histograms of logarithmic 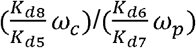 **(F)** scatter plot of magnitude of Bell curves, defined as the fold-increase of peak *EE* from the baseline level when *X*=0, vs. logarithmic 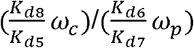.

There is only a small overlap between the two, as indicated by the histograms of 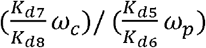 of the two curve types (Fig. 9B). Moreover, the magnitude of the J/U curves, defined as the fractional-decrease of the nadir *EE* from the baseline level when *X*=0, is positively associated with 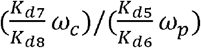 (Fig. 9C). These results are consistent with condition C1 postulated for the emergence of J/U curves.

Among the MD and Bell curves, the paired values of 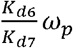 and 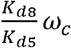 are distributed predominantly above the diagonal for Bell curves, and below the diagonal for MD curves (Fig. 9D). There is a small overlap between the two, as indicated by the histogram of 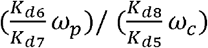 (Fig. 9E). Moreover, the magnitude of the Bell curves, defined as the fold-increase of the peak *EE* from the baseline level when *X*=0, is positively associated with 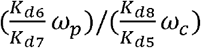 (Fig. 9F). These results are consistent with condition C2 postulated for the emergence of Bell curves.

Examining the individual parameters associated with the 4 curve shapes revealed distribution patterns that are consistent with conditions C1 and C2 for differentiating these curves (Fig. 10). For the MI vs. J/U curves, although the distributions of each parameter substantially overlapped between the two curve types, they are biased in directions that favor C1. For the MD vs. Bell curves, although the biases in the *K*_*d*5_, *K*_*d*6_, *K*_*d*7_, *K*_*d*8_ distributions are not as pronounced as those for the MI vs. J/U curves situation, they still have tendencies that favor C2 with *K*_*d*6_ and *K*_*d*8_ distributions being the least biased. In contrast, *ω*_*p*_ and *ω*_*c*_ are predominantly in small values for MD and Bell curves respectively (Fig. 10E and 10F), a pattern that is also consistent with favoring C2.

**Figure 10.**
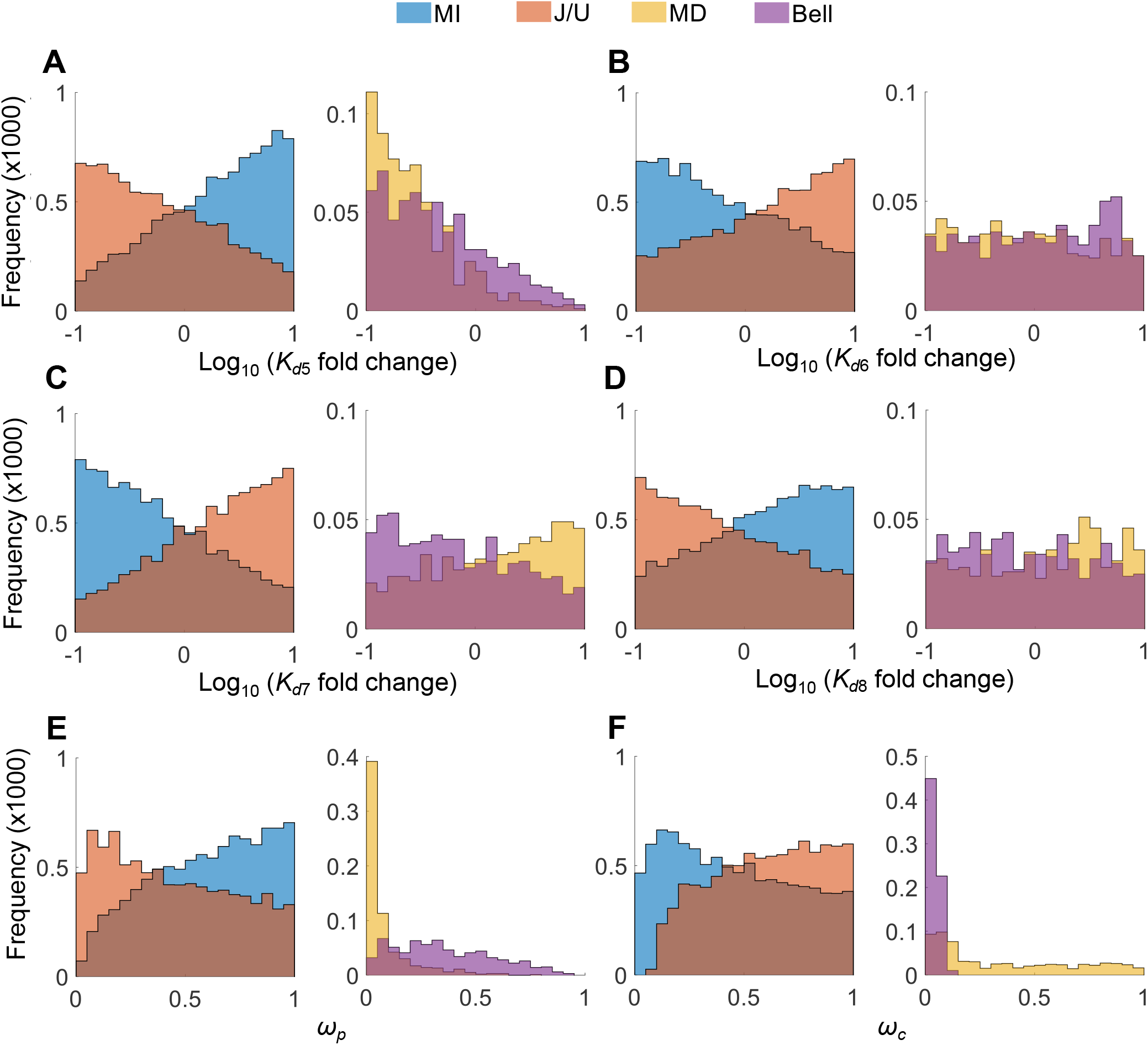
Distributions of individual parameters as indicated for MI vs. J/U and MD vs. Bell curves from the 20,000 six-parameter MC simulations as presented in Fig. 9 and S8.

#### 3.2 Population MC simulations

The above MC simulations were limited to 6 key parameters while all other parameters remained constant. To validate the relationships between these parameters for nonmonotonic endocrine effect in a more unbiased way, we next conducted MC simulations by using the virtual population HPE model where each individual has different *EH* and *PH* levels which are determined by varying values of all relevant model parameters as detailed in Methods. The population MC simulations generate a variety of shapes of DR curves, which include, in addition to MI, J/U, MD, and Bell, also multi-phasic curves, i.e., U-then-Bell and Bell-then-U shapes (Fig. S9A-S9F). The percentage distributions of these different shapes are 56, 28, 9, 4, 0.04, and 3% respectively (Fig. S9G). Therefore, MI and MD curves together are nearly twice as frequent as NMDR curves.

Similar to the six-parameter MC simulations above, the population MC simulations showed that conditions C1 and C2 are largely observed for MI vs. J/U (Fig. 11A-11B) and MD vs. Bell (Fig. 11D-11E) curves respectively, although the separations are not as clean. The magnitudes of the J/U and Bell curves are positively associated with 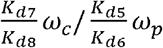 (Fig. 11C) and 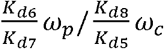 (Fig. 1F), respectively. Examining the 6 individual parameters *K*_*d*5_, *K*_*d*6_, *K*_*d*7_, *K*_*d*8_, *ω*_*p*_, and *ω*_*c*_ for the 4 curve shapes revealed distribution patterns that are largely consistent with conditions C1 and C2 to differentiate these curves (Fig. 12A-12F), with distribution biases qualitatively similar to the six-parameter MC simulations above (Fig. 10). For the remaining parameters, *k*_1_, *k*_30_, *k*_3_, *K*_d3_, *n*_3_, and *CR*_*tot*_, there are some interesting distribution patterns. The distributions of *k*_1_ are largely indistinguishable among the 4 curve chapes (Fig. 12G) and so are those of *n*_3_ (Fig. 12K). In contrast, *k*_30_ and *k*_3_ tend to have high and low values respectively to favor monotonic DR curves (Fig. 12H and 12I). For *K*_*d*3_, the differences in the distributions for monotonic vs nonmonotonic curves are small, with a slight tendence of lower values favoring J/U curves (Fig. 12J). The differences in the distributions of *CR*_*tot*_ for monotonic vs nonmonotonic curves are also small, with a slight tendence of higher values favoring J/U curves (Fig. 12L).

**Figure 11.**
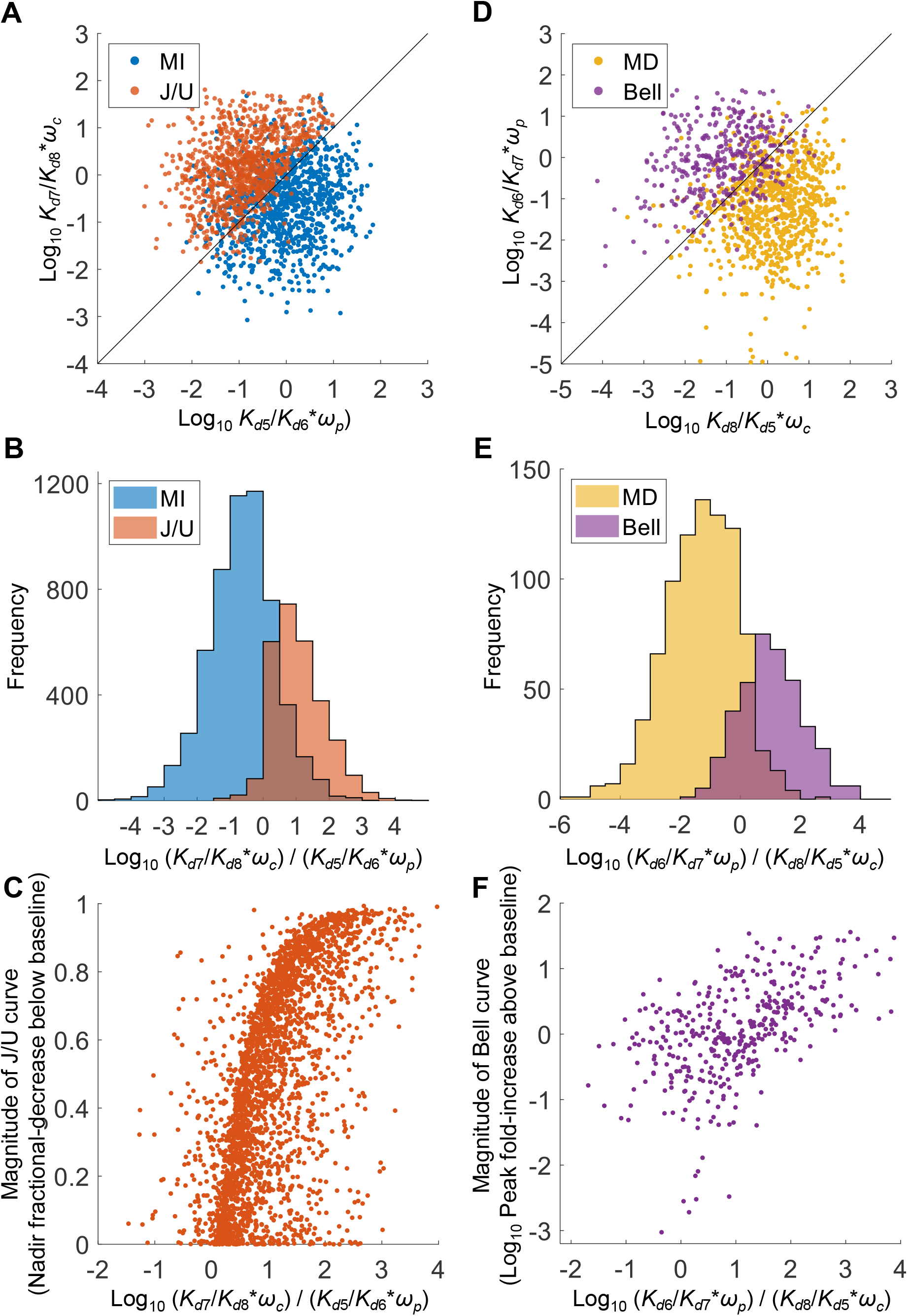
Relationships between *K*_*d*5_, *K*_*d*6_, *K*_*d*7_, *K*_*d*8_, *ω*_*p*_, and *ω*_*c*_ for MI vs. J/U and MD vs. Bell curves from population MC simulations of 9,996 individuals. The parameters *k*_5_, *k*_6_, and *k*_8_ were randomly sampled from uniform distributions of log_10_([0.1, 10]) as fold change relative to the respective default values, and *ω*_*p*_ and *ω*_*c*_ were randomly sampled from the uniform distribution [0,1]. For MI and J/U curves, **(A)** scatter plot of logarithmic 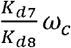 vs. 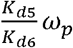 of 1000 randomly selected paired values, **(B)** distribution histograms of logarithmic 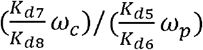, **(C)** scatter plot of magnitude of J/U curves vs. logarithmic 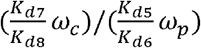. For MD and Bell curves, **(D)** scatter plot of logarithmic 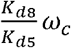 vs.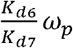, **(E)** distribution histograms of logarithmic 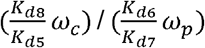, **(F)** scatter plot of logarithmic magnitude of Bell curves vs. logarithmic 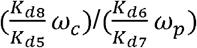.

**Figure 12.**
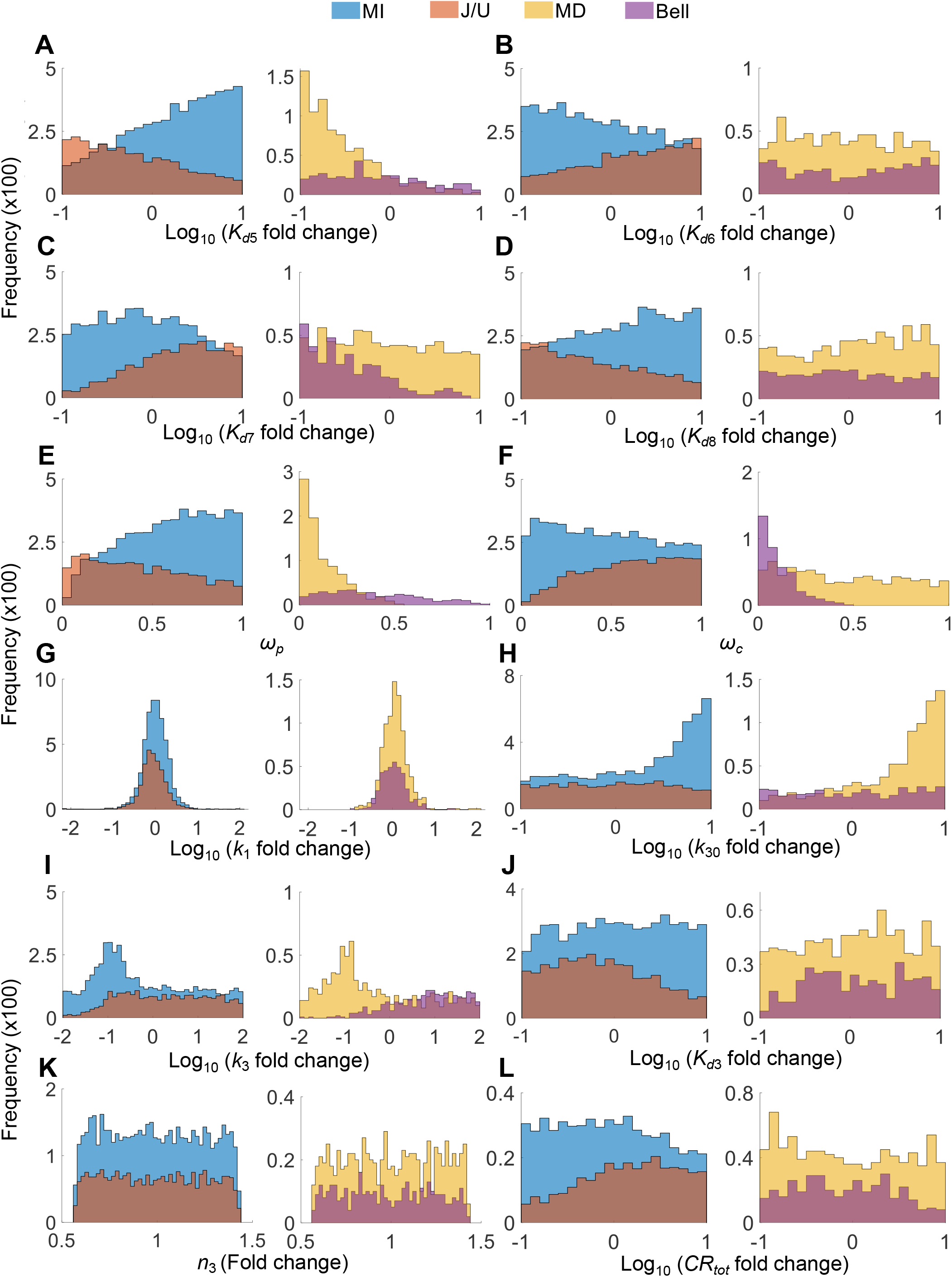
Distributions of parameters as indicated for MI vs. J/U and MD vs. Bell curves from the population MC simulations of 9,996 individuals as presented in Fig. 11 and S9.

## Discussion

In the present study we demonstrated that negative feedback regulation, common and intrinsic to nearly all homeostatic endocrine systems, are theoretically capable of rendering nonmonotonic responses to EDC perturbations. In essence, when an EDC is able to sufficiently interfere with the central feedback action to affect the pituitary hormone and in turn the endogenous effector hormone levels, it can produce at low doses a net endocrine effect that is in an opposite direction of what is normally expected for an agonist or antagonist. For NMDR to arise, we showed for the first time that certain parameter conditions must be met, as indicated in C1 and C2. These conditions require that the effector hormone and EDC have differential binding affinities and efficacies for the peripheral target receptor that mediates the endocrine effect and the central receptor that mediates the negative feedback regulation. Provided C1 or C2 are met, other parameters such as *k*_*30*_ and *k*_*3*_ which govern PH synthesis may have a modulatory role, enhancing or attenuating the NMDR magnitude. For the J/U-shaped response, it requires that an EDC agonist has a stronger inhibitory effect in the central negative feedback pathway than its stimulatory effect in the peripheral target tissues. In this case the endogenous hormone would be sufficiently downregulated at low EDC concentrations, resulting in an overall reduction in the endocrine effect despite that locally the EDC is an agonist in the peripheral tissue. For the Bell-shaped response, it requires that an EDC antagonist has a stronger action to block the endogenous effector hormone-mediated central negative feedback than its inhibitory action in the peripheral target tissues. In this case, the endogenous hormone would be upregulated at low EDC concentrations, resulting in an overall increase in the endocrine effect despite that locally the EDC is an antagonist in the peripheral tissue. In both cases, at high concentrations when the central action of the EDC is saturated, its peripheral action will take over, reversing the direction of the endocrine effect exhibited at low concentrations.

### 1. Nonmonotonicity via incoherent feedforward action

The NMDRs predicted by the HPE axis model here are essentially a result of incoherent feedforward actions of EDCs (Kaplan, Bren et al. 2008, Zhang, Pi et al. 2009). In this framework (Fig. 1B), an EDC acts in two opposing arms: it has (i) a direct endocrine effect in the target issue, and (ii) an indirect but opposite endocrine effect by altering the endogenous effector hormone level through interfering with the HPE feedback. The EDC concentration-dependent changes in the signaling strengths of the two arms determine the change in the direction of the endocrine effect and the shape of the DR curves. Through signal amplification in the hypothalamus and pituitary, the indirect arm can be perturbed by the EDC to readily alter the endogenous hormone levels, leading to “overcorrection” of the endocrine effect exerted by the EDC via the direct arm. In this regard, the higher the feedback amplification gain, the smaller the differences are required of the EDC*’*s relative binding affinities and efficacies between the central and peripheral actions to produce NMDRs.

### 2. Agonistic vs. antagonistic actions

For a given receptor-mediated biological effect in a tissue, the efficacy *ω* of an EDC, relative to the background action level of the endogenous hormone, determines whether it is a (partial) agonist or antagonist in that endocrine context (Howard and Webster 2009, Howard, Schlezinger et al. 2010). In the current model specifically, with the receptor occupancy of both *PR* and *CR* by the endogenous effector hormone at 10% as the average baseline, whether *ω*_*p*_ and *ω*_*c*_ are greater or smaller than 0.1 largely determines whether the local action of the EDC is agonistic or antagonistic at the peripheral and central sites, respectively. A closer examination of the distributions of *ω*_*p*_ and *ω*_*c*_ for all 4 types of DR curves revealed that this indeed seems to be the case (Fig. S10). The MI and MD curves, representing mainly agonistic and antagonistic actions of the EDC in the peripheral target tissue respectively, can be largely distinguished by *ω*_*p*_ levels. For the six-parameter MC simulations, *ω*_*p*_ is highly concentrated in the range of 0.1-1 for MI curves, whereas it is mostly < 0.1 for MD curves (Fig. S10A). Similar dichotomy was found with the population MC simulations (Fig. S10E). In comparison, *ω*_*c*_ can vary in the entire range of 0-1 regardless of MI or MD curves (Fig. S10B and S10F). Therefore, the peripheral agonistic or antagonistic action of the EDC relative to the local endogenous effector hormone level there seems to play a primary role in producing MI or MD curves.

In contrast, J/U vs. Bell curves can be largely distinguished by *ω*_*c*_ levels. For the six-parameter MC simulations, *ω*_*c*_ is highly concentrated in the range of 0.1-1 for J/U curves, whereas it is mostly < 0.1 for Bell curves (Fig. S10D). A similar dichotomy was found with the population MC simulations, albeit the separation is not as clean (Fig. S10H). In comparison, *ω*_*p*_ can vary in the entire range of 0-1 regardless of J/U or Bell curves (Fig. S10C and S10G). Therefore, the central agonistic or antagonistic action of the EDC relative to the local endogenous effector hormone level there seems to play a primary role in producing one of the two types of NMDRs. Similar conclusions can be drawn when the baseline receptor occupancy is considerably higher (e.g., 50%) or lower (e.g., 1%) than the default 10% (simulation results not shown).

### 3. Selective receptor modulators and complex NMDRs

The binding affinity of the endogenous hormone or an exogenous compound for the hormone receptor may vary in different cells and tissues, depending on the status of posttranslational covalent modifications such as phosphorylation, oxidation, acetylation, and methylation, and the intracellular milieu (Faus and Haendler 2006,Malbeteau, Pham et al. 2021). Upon ligand binding, downstream molecular events, such as receptor dimerization, DNA binding, and co-regulator recruitment, can determine the efficacy, thus the direction and magnitude of the endocrine effect. Variations in these molecular events can cause differential binding affinities and efficacies, which may lead to NMDRs for different endocrine active compounds in different tissues. These variations may contribute to the phenomenon of tissue-specific selective receptor modulators (SRM) for estrogen, progesterone, androgen, and thyroid hormone receptors (Riggs and Hartmann 2003, Christiansen, Lipshultz et al. 2019, Islam, Afrin et al. 2020, Saponaro, Sestito et al. 2020). In keeping with this concept, our modeling showed that an EDC can be stimulatory in some endocrine contexts while inhibitory in others. For environmental exposures which often involve mixtures of EDCs, the direction of the endocrine effects will be ultimately determined by the net actions of different compounds possessing different binding affinities and efficacies acting potentially at the central and/or peripheral sites simultaneously. Some constituents in a mixture may act primarily at the peripheral site while others may act primarily at central site, thus creating complex endocrine outcome scenarios. Genetic and epigenetic variations between human individuals may also result in different binding affinities and efficacies in central and peripheral tissues even for the same compound, which may lead to emergence of NMDR only in certain subpopulations, as suggested by our MC population simulations. Lastly, in women whose circulating estradiol and progesterone levels fluctuate through the menstrual cycle, the net endocrine effect of an EDC may vary depending on the phase of the cycle. The present study suggests that the DR relationship induced by EDCs can be more complex than J/U and Bell-shape. In the MC population simulation, there are cases where an EDC exhibits U-then-Bell or Bell-then-U curves (Fig. S9E and S9F). Whether such complex responses occur *in vivo* remains to be determined.

### 4. Feedback mechanisms proposed for NMDR in the literature

Negative feedback has been frequently referred to in the EDC literature as one of the underlying mechanisms for NMDR (Vandenberg, Colborn et al. 2012, Lagarde, Beausoleil et al. 2015), yet there are barely any studies that have provided evidence or rigorous arguments on how NMDR may arise in this context, at least in theory. In the review article (Vandenberg, Colborn et al. 2012), negative feedback control in endocrine systems such as insulin-glucose and TSH-TH was proposed as an NMDR-producing mechanism, however the studies cited were concerned with temporal responses of sexual organ growth to steroid hormone stimulation without reporting dose-responses. Specifically, these studies described a plateauing of the proliferative growth of the prostate gland stimulated by androgen (Lesser and Bruchovsky 1974, Bruchovsky, Lesser et al. 1975), a similar plateau response of uterus growth to estrogen (Wiklund, Wertz et al. 1981), or refractory uterine cell proliferation after successive estrogen treatments (Stormshak, Leake et al. 1976). The lack of further growth of these organs was interpreted as a result of engagement of some negative feedback mechanisms that eventually limit cell proliferation. In the review article (Lagarde, Beausoleil et al. 2015), a number of diverse studies were cited to support negative feedback as a potential mechanism for NMDR. Major NMDR findings in these studies include Na^+^/H^+^ exchanger activity in response to 17β-estradiol (E2) in rat aortic smooth muscle cells (Incerpi, D’Arezzo et al. 2003), puberty onset in rats in response to BPA (Adewale, Jefferson et al. 2009), transcriptional induction of hepatic lipogenic genes in mice by BPA (Marmugi, Ducheix et al. 2012), enhancement of spatial memory in rats by E2 (Inagaki, Gautreaux et al. 2010), mouse mammary growth in response to diethylstilbestrol (DES) or E2 (Skarda 2002, Skarda 2002, Köhlerová and Skarda 2004, Vandenberg, Wadia et al. 2006), and mouse prostate enlargement by fetal exposure to E2 or DES (vom Saal, Timms et al. 1997). However, in none of these studies was the global negative feedback, as we explored here, explicitly discussed as a mechanism of NMDR. Rather, they suggested that hormone receptor desensitization or downregulation, occurring locally in cells, is a potential mechanism. Whether receptor downregulation itself is a result of intracellular negative feedback upon receptor activation is another topic, and even so, only conditional not steady-state NMDR, as we explored here, is expected (Zhang, Pi et al. 2009). Taken together, compared with past attempts in this area, our modeling study here revealed, for the first time, the mechanistic rationale and parameter conditions by which NMDR may arise through interference of EDCs with the systemic negative feedback action of the endogenous hormones.

### 5. Implications and potential applications in risk assessment of EDCs

The NMDR effect predicted by our model suggests that at low concentrations the endocrine outcome of an EDC *in vivo* may run in the opposite direction of what is expected based on findings from unsophisticated *in vitro* bioassays, such as receptor binding and receptor-mediated reporter assays. With the advent of new approach methodologies (NAM), this possibility of counterintuitive effects will pose a serious challenge to the task of *in vitro* to *in vivo* extrapolation (IVIVE), where *in vitro* derived point-of-departure (PoD) concentrations are used to extrapolate human reference doses for EDC safety regulation. To circumvent such potential *in vitro* to *in vivo* discordance, developing sophisticated endocrine-system-on-a-chip – by incorporating multiple interacting organoids or cell cultures to mimic the global negative feedback structure as *in vivo* – could be a solution moving forward. However, technical difficulties aside, covering sufficient space of population variability with such endocrine-system-on-a-chip assays to predict nonmonotonic effects for susceptible subpopulations will pose another daunting challenge.

Alternatively, developing quantitative adverse outcome pathway (qAOP) models of endocrine systems as we initiated here may be a viable computational approach to aid the IVIVE task for EDCs. In the qAOP models, the inter-individual differences in the synthesis, secretion, metabolism, and actions of the hormones can be appropriately coded in the values of nominal parameters associated with these processes to account for individual responses. These population models will allow us to help identify the risk factors for NMDRs and susceptible subpopulations.

### 6. Limitations and future directions

The criteria of C1 and C2 for the emergence of NMDRs are based on the assumption that the EDC concentrations in the brain and peripheral target tissue are at comparable levels. In reality, due to the existence of the blood brain barrier (BBB) and membrane transporters, the partitioning of an EDC into the brain matter such as the hypothalamus can be quite different than in the peripheral tissues, resulting in either lower or higher central concentrations (Denuzière and Ghersi-Egea 2022). One exception is the anterior pituitary, which is a circumventricular organ sitting outside of the BBB (Ganong 2000, Kiecker 2018), and therefore can be readily accessed by circulating EDCs. Similarly, differential central vs. peripheral partitioning also applies to endogenous hormones (Martin, Plank et al. 2019, Colldén, Nilsson et al. 2022). As a result, C1 and C2 should be revised to account for the differential site concentrations as follows:

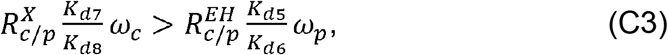

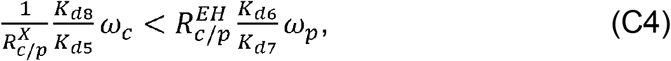

where 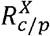 is the concentration ratio of the central to peripheral EDC, and 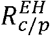 the concentration ratio of the central to peripheral endogenous effector hormone. Many endocrine active compounds have dual-site actions including pharmaceutical drugs. For instance, the thyromimetic drug sobetirome and its prodrug Sob-AM2 can penetrate easily into the CNS to inhibit TRH and TSHβ expression, which causes a severe depletion of circulating T4 and T3 in rats (Ferrara, Bourdette et al. 2018). However, animals treated with these drugs remained free of clinical signs of hypothyroidism, due to the thyromimetic actions of the drugs in thyroid hormone-target tissues.

The majority of the steroid hormones in the blood circulation exist in bound forms in complex with binding proteins (Hammond 2016), a process not considered in the current model. Since we deal with steady-state responses and it is the free hormones, whose steady-state levels are only determined by the HPE feedback loop, that move into the central and target tissues to take action (Bikle 2021), excluding the partitioning with the plasma proteins does not affect the DR curves and the findings of the present study.

In the present study, we only considered the scenario of an EDC antagonist acting in passive mode, where the EDC competes with the endogenous ligand for the receptors and the efficacy *ω* can drop to as low as zero. It is also possible that an EDC may act as an active antagonist by recruiting co-repressors to downregulate the basal ligand-independent transcriptional activities of the target genes (Stoney Simons 2003, Heldring Pawson et al. 2007). In such situations, *ω* may become negative. However, the overall conclusions are not expected to change, because condition C2 and C4 can still be readily met to achieve Bell-shaped DR.

In the current model, the peak or nadir of an NMDR curve of the endocrine effect is associated with a considerable change in the endogenous hormone level. This may occur with pharmaceutical compounds such as sobetirome and Sob-AM2 that can nearly deplete thyroid hormones in rats (Ferrara, Bourdette et al. 2018). However, if the entry of the EDC to the brain is mediated by a transporter-mediated saturable process, the central action of the EDC to interfere with the HPE feedback may be capped within a limit. When this happens, the inflection point of the NMDR curve may occur at only moderately altered endogenous hormone levels.

In the present study we used the HPE feedback framework as a generic example to investigate the biological mechanisms and conditions for NMDRs. The mechanistic principles identified here should be applicable also to other endocrine feedback systems not explicitly involving the hypothalamus and pituitary. Examples are feedback regulations between insulin and glucose for blood sugar control, and between parathyroid hormone, VD3, and calcium for calcium homeostasis. Future work may customize the generic HPE feedback model toward specific endocrine systems and explore the corresponding biological conditions for NMDRs.

## Conclusions

In summary, through systems modeling the present study revealed a potentially universal mechanism for the emergence of NMDRs frequently encountered with EDCs. By interfering with the systemic negative feedback action of the endogenous hormones, EDCs may present counterintuitive low-dose effects and NMDRs. These nontraditional DR behaviors emerge when an EDC has differential receptor binding affinities and efficacies relative to the endogenous hormones in the central and peripheral tissues. Through populational simulations, our modeling provided novel insights into the inter-individual variabilities in response to EDC exposures and future studies may help identify risk factors and susceptible subpopulations for health safety assessment and protection.

## Funding Acknowledgements

This research was supported in part by the National Institute of Environmental Health Sciences grants P42ES04911 and R01ES032144 and Department of Defense grant HT9425-23-1-0809.

## Author Contributions

**ZS:** Formal analysis, Investigation, Methodology, Writing – original draft, Writing – review & editing, Visualization. **SX:** Funding acquisition, Writing – review & editing. **QZ:** Conceptualization, Formal analysis, Investigation, Methodology, Funding acquisition, Project administration, Supervision, Validation, Writing – original draft, Writing – review & editing.

## Notes

**Conflict of Interest:** All authors declare no conflict of interest.

### Competing Interest Statement

The authors have declared no competing interest.

