## Supplemental Material for "Interference with Systemic Negative Feedback Regulation as a Potential Mechanism for Nonmonotonic Dose-Responses of Endocrine-Disrupting Chemicals"

### Processing of thyroid profile dataset in NHANES (2007-2012)

Records in all three cycles were resampled based on the assigned sample weights to correct for oversampling of subpopulations in the NHANES data. In each cycle, each record's (individual's) sample weight was first normalized by dividing with the maximum weight in that cycle. Based on the normalized sample weights, a rejection sampling method was applied to determine whether a record is selected into a final weight-adjusted population. For simplicity, a uniform distribution unif(0, 1) was used as the candidate distribution [1]. The higher the sample weight of an individual, the more likely the individual will be selected. In this case, a total of 2172 records were selected. Furthermore, individuals who have taken thyroid drugs and had thyroid cancer were excluded, resulting in 2035 records in a weight-adjusted population. Lastly, records with missing fT4 or TSH values were removed, resulting in 1883 individuals in a final weight-adjusted population.

**Table S1.** Model ODEs

| | | |
|---|---|---|
| $\frac{dEH}{dt}$ | = | $k_1 \cdot PH - k_2 \cdot EH$ |
| $\frac{dPH}{dt}$ | = | $k_{30} + \frac{k_3 \cdot {K_{d3}}^{n_3}}{{K_{d3}}^{n_3} + (EHCR + \omega_c \cdot XCR)^{n_3}} - k_4 \cdot PH$ |
| $\frac{dEHCR}{dt}$ | = | $k_{f7} \cdot EH \cdot CR - k_{b7} \cdot EHCR$ |
| $\frac{dXCR}{dt}$ | = | $k_{f8} \cdot X \cdot CR - k_{b8} \cdot XCR$ |
| $\frac{dEHPR}{dt}$ | = | $k_{f5} \cdot EH \cdot PR - k_{b5} \cdot EHPR$ |
| $\frac{dXPR}{dt}$ | = | $k_{f6} \cdot X \cdot PR - k_{b6} \cdot XPR$ |

**Table S2**. Model parameters and default values

| Parameter | Default value | Unit | Note |
|---|---|---|---|
| $k_1$ | 0.2888 | Conc/h | Rate constant of *PH*-stimulated *EH* production. Together with the $k_2$ value below, the default value produces a steady-state concentration of *EH* at 10. |
| $k_2$ | 0.0289 | 1/h | Rate constant of *EH* degradation. The default value corresponds to a half-life of 24 h. |
| $k_{30}$ | 0.1 | Conc/h | Basal *PH* production rate, which allows the minimally |

| | | | |
|---|---|---|---|
| | | | reachable *PH* level upon full negative feedback inhibition to be 10-fold below the baseline *PH* level. |
| $k_3$ | 10 | Conc/h | Maximal *PH* production rate, which allows the maximally reachable *PH* level upon no negative feedback inhibition to be near 10-fold above the baseline *PH* level. |
| $K_{d3}$ | $\sqrt[7]{0.09/0.91}$ | Conc | Affinity constant of *CR*-mediated negative feedback inhibition of *PH*. The default value is set to make sure that the baseline production of *PH* is 10% of the maximum. |
| $n_3$ | 7 | - | Hill coefficient. The default value is set to 7 to describe the ultrasensitivity of the negative feedback loop, which is a reasonable estimate based on feedback gain of the hypothalamic-pituitary-thyroid axis [2-4]. |
| $k_4$ | 1 | 1/h | Rate constant of *PH* degradation. The default value corresponds to a half-life of 0.693 h. |
| $K_{d5}$ | 90 | Conc | $K_{d5} = k_{b5} / k_{f5}$. Dissociation constant of the reversible binding of *EH* + *PR* ↔ *EHPR*. The default value produces 10% *PR* occupancy at the baseline *EH* level. |
| $k_{f5}$ | 1 | 1/Conc/h | Association rate constant. |
| $k_{b5}$ | 90 | 1/h | Dissociation rate constant. The default value corresponds to a 40-second mean residence time of *EH* on *PR*. Since we are dealing with steady-state responses, the actual value here does not matter. |
| $K_{d6}$ | 90 | Conc | $K_{d6} = k_{b6} / k_{f6}$. Dissociation constant of the reversible binding of *X* + *PR* ↔ *XPR*. The default value is set to be the same as $K_{d5}$. |
| $k_{f6}$ | 1 | 1/Conc/h | Association rate constant. Default value set same as $k_{f5}$. |
| $k_{b6}$ | 90 | 1/h | Dissociation rate constant. Default value set same as $k_{b5}$. |
| $K_{d7}$ | 90 | Conc | $K_{d7} = k_{b7} / k_{f7}$. Dissociation constant of the reversible binding of *EH* + *CR* ↔ *EHCR*. The default value produces 10% *CR* occupancy at the baseline *EH* level. |
| $k_{f7}$ | 1 | 1/Conc/h | Association rate constant. |
| $k_{b7}$ | 90 | 1/h | Dissociation rate constant. The default value corresponds to a 40-second mean residence time of *EH* on *CR*. Since we are dealing with steady-state responses, the actual value here does not matter. |
| $K_{d8}$ | 90 | Conc | $K_{d8} = k_{b8} / k_{f8}$. Dissociation constant of the reversible binding of *X* + *CR* ↔ *XCR*. The default value is set to be the same as $K_{d7}$. |
| $k_{f8}$ | 1 | 1/Conc/h | Association rate constant. Default value set same as $k_{f7}$. |
| $k_{b8}$ | 90 | 1/h | Dissociation rate constant. Default value set same as $k_{b7}$. |
| $w_p$ | 1 (agonist)<br>0 (antagonist) | - | Efficacy of *X* for *PR* |
| $w_c$ | 1 (agonist) | - | Efficacy of *X* for *CR* |

| | | | |
|---|---|---|---|
| | 0 (antagonist) | | |
| $PR_{tot}$ | 10 | Conc | Total concentration of peripheral receptors |
| $CR_{tot}$ | 10 | Conc | Total concentration of central receptors |

Abbreviations: Conc: concentration, h: hour.

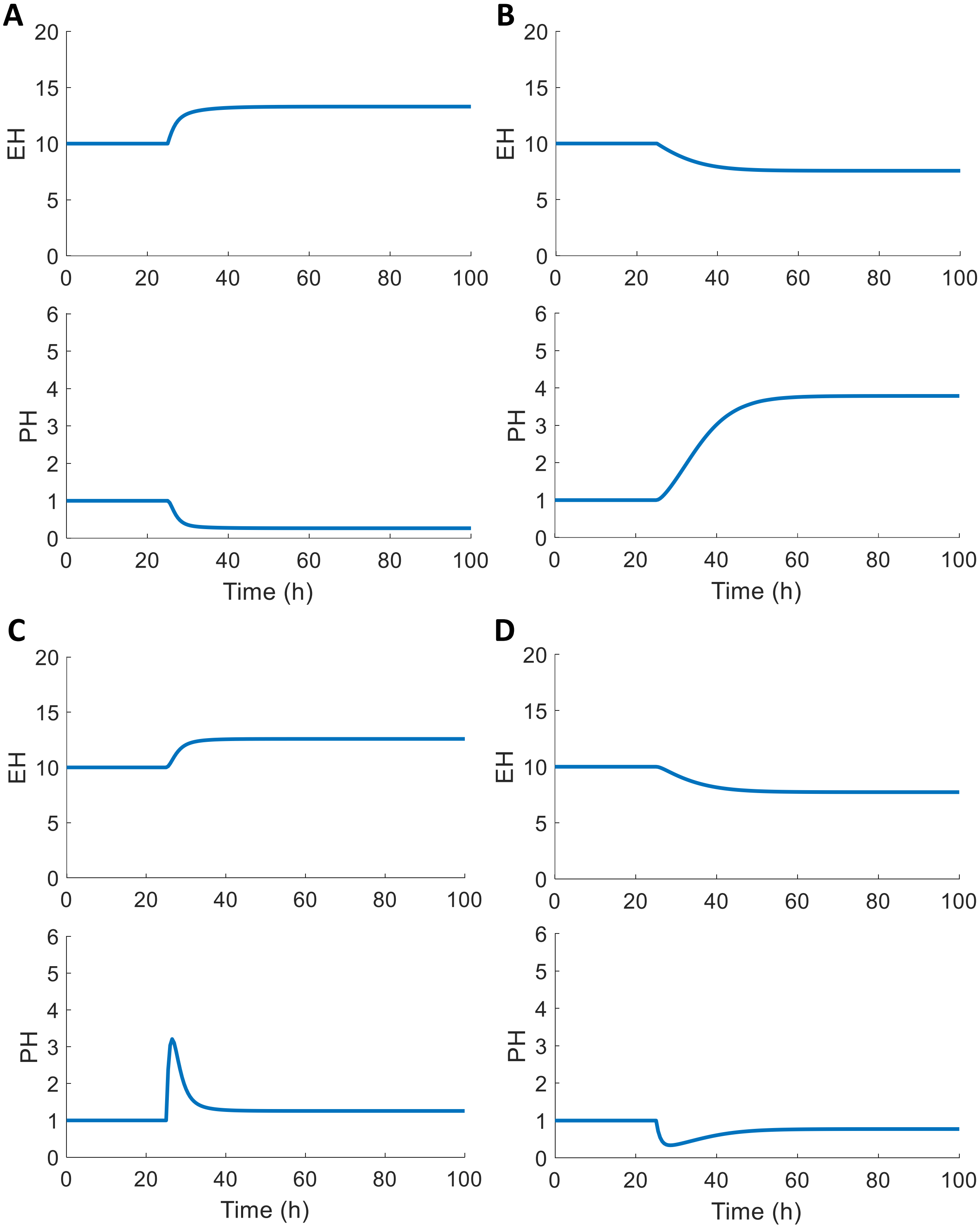


**Figure S1.** Dynamic responses of *EH* and *PH* simulating primary and secondary endocrine disorders. **(A)** Primary hyper-functioning condition by increasing $k_1$ by 5-fold. **(B)** Primary hypo-functioning condition by decreasing $k_1$ by 5-fold. **(C)** Secondary hyper-functioning condition by increasing $k_3$ by 5-fold. **(D)** Secondary hypo-functioning condition by decreasing $k_3$ by 5-fold.

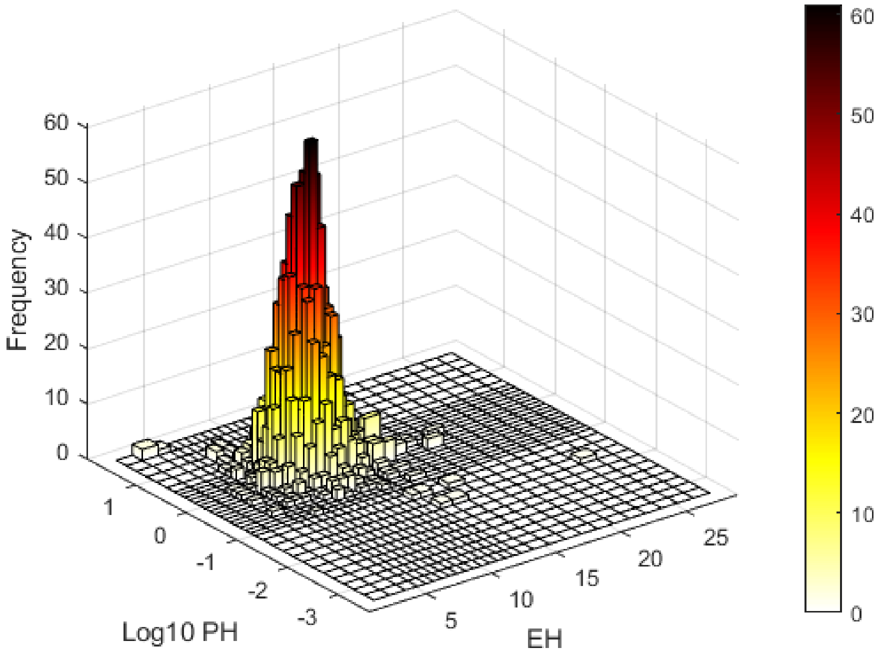


**Figure S2.** Population density of *PH* and *EH* levels based on the respective TSH and fT4 concentrations in 2007-2012 NHANES thyroid profile dataset.

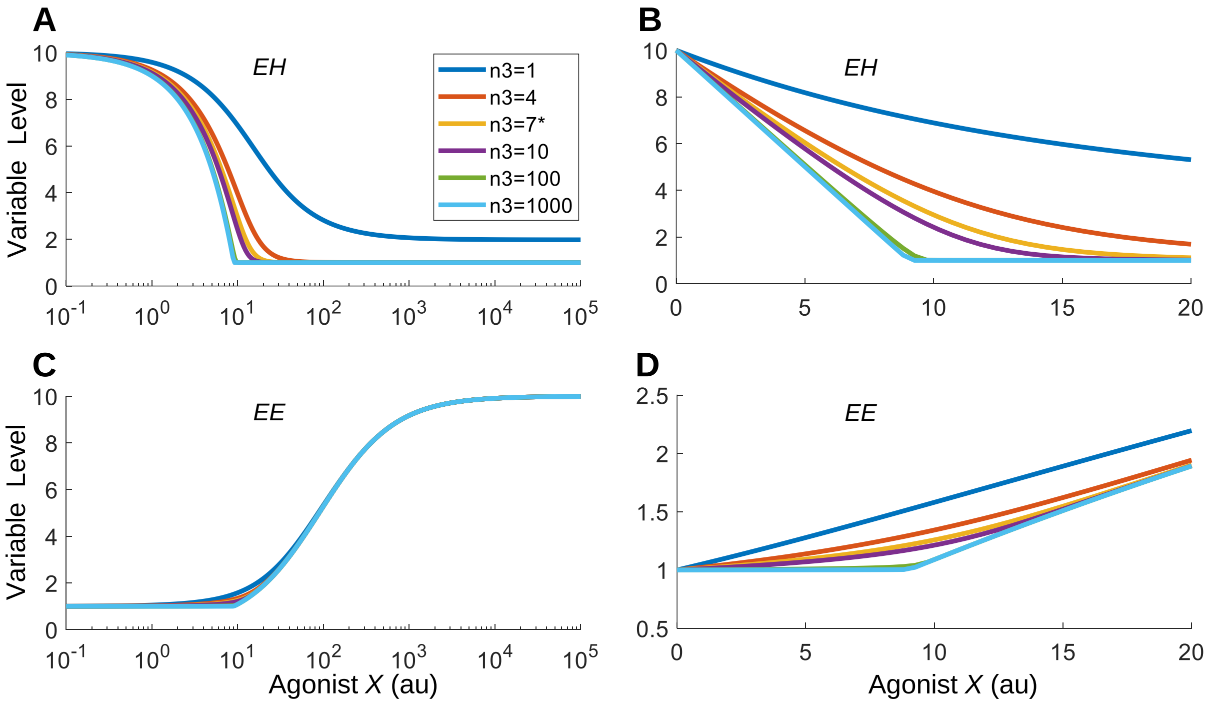


**Figure S3.** The effect of feedback gain (Hill coefficient $n_3$) on the steady-state DR profiles for variables as indicated when *X* acts as the reference agonist with identical binding affinities and efficacies as *EH* for *CR* and *PR*. (A-B) The *EH* vs. *X* DR on $\log_{10}$ and linear x-axis, respectively. (C-D) The *EE* vs. *X* DR on $\log_{10}$ and linear x-axis, respectively. The color-coding scheme is indicated in (A) for different values of $n_3$. * denotes default $n_3$ value. With high $n_3$ values, the *EH* vs. *X* DR curve is linearized and decreases at low *X* concentrations (B), and *EE* is flat at low *X* concentrations (C and D).

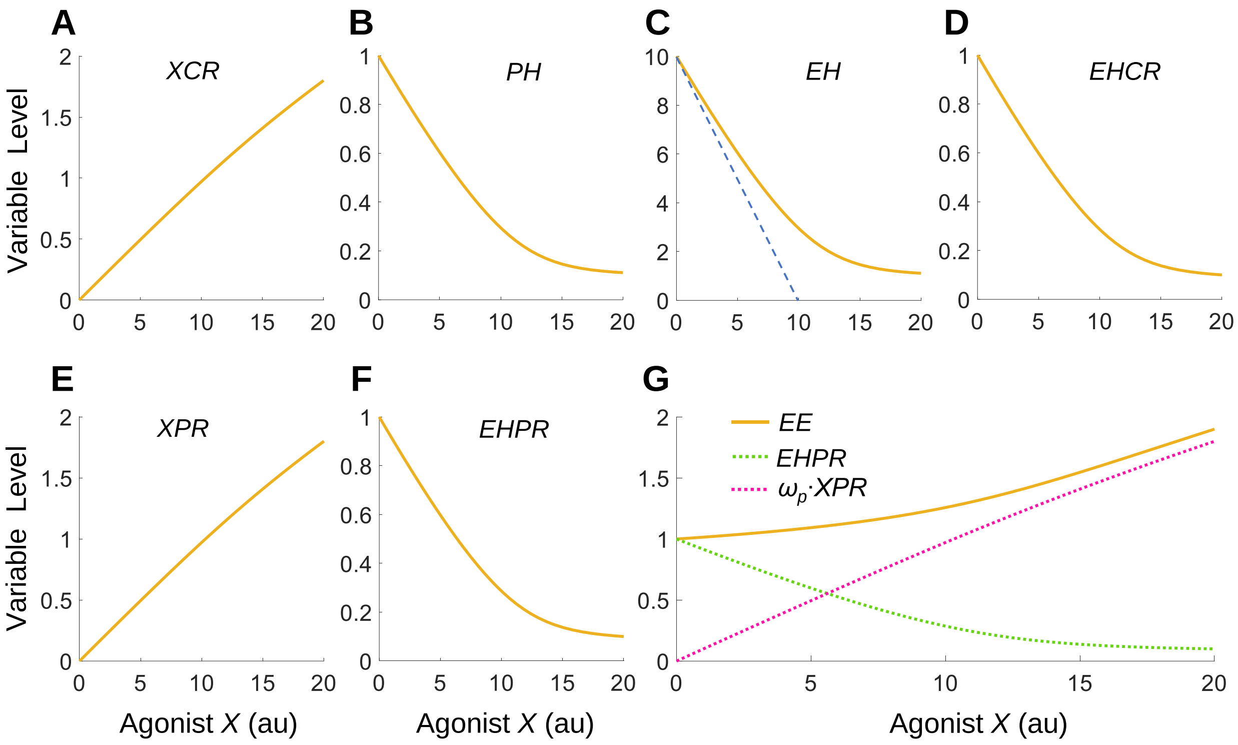


**Figure S4.** Same results as in Fig. 2 except that the x-axis is in linear scale and spans a much smaller low-level $X$ range. Blue dash line in (C) denotes the case of linearization when $n_3$ is infinitely high.

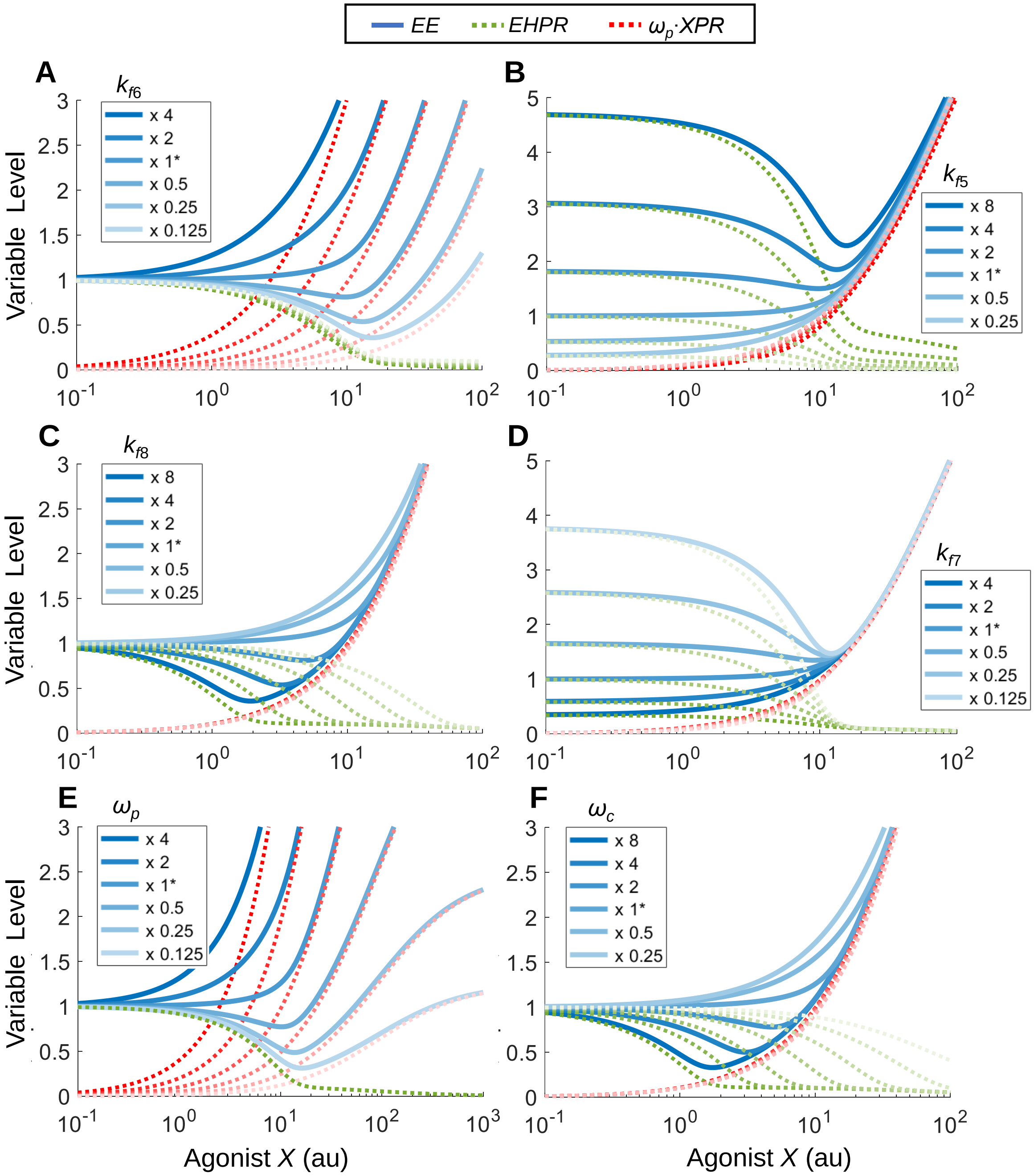


**Figure S5.** Illustration of the mechanisms of J/U-shaped DR of *EE* in response to agonist *X* when parameters $k_{f6}$ (A), $k_{f5}$ (B). $k_{f8}$ (C), $k_{f7}$ (D), $\omega_p$ (E), and $\omega_c$ (F) are varied. x 1* denotes the parameter is at default value, and x 0.125, 0.25, x 0.5, x 2, x 4, and x 8 denote that the parameter is at corresponding fold of the default value. Lighter shade of the curves is associated with lower parameter values as indicated. Same denotation is used in other figures where applicable.

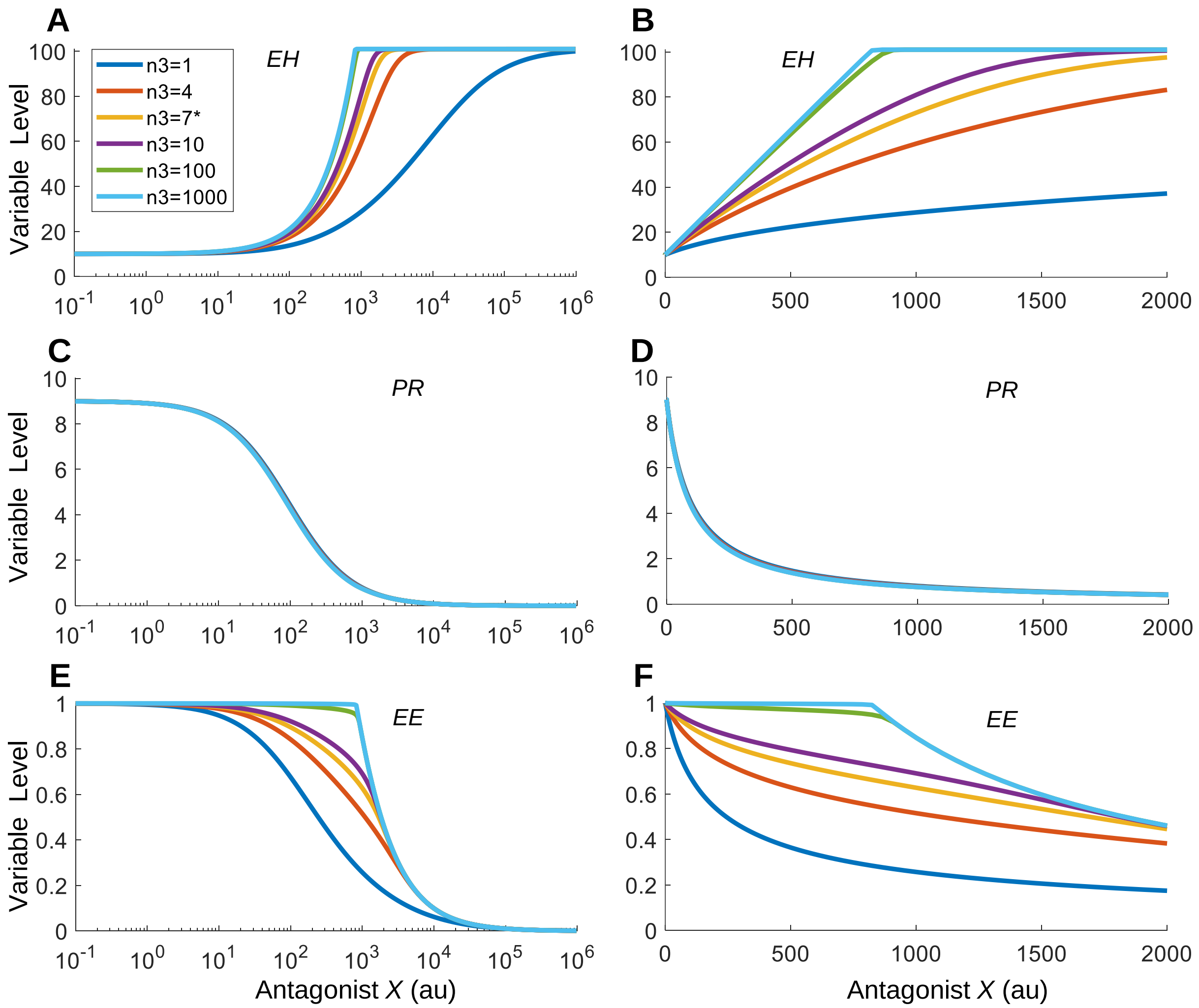


**Figure S6.** The effect of feedback Hill coefficient $n_3$ on the steady-state DR profiles for variables as indicated when $X$ acts as the reference antagonist with identical binding affinities as the endogenous hormone *EH* for *CR* and *PR* but the efficacies are zero. (A-B) The *EH* vs. $X$ DR on $\log_{10}$ and linear x-axis, respectively. (C-D) The *PR* vs. $X$ DR on $\log_{10}$ and linear x-axis, respectively. (E-F) The *EE* vs. $X$ DR on $\log_{10}$ and linear x-axis, respectively. Color-coding scheme is indicated in (A) for different values of $n_3$. * denotes default $n_3$ value. With high $n_3$ values, the *EH* vs. $X$ DR curve is linearized and increases at low $X$ concentrations (B) and *EE* is flat (E and F).

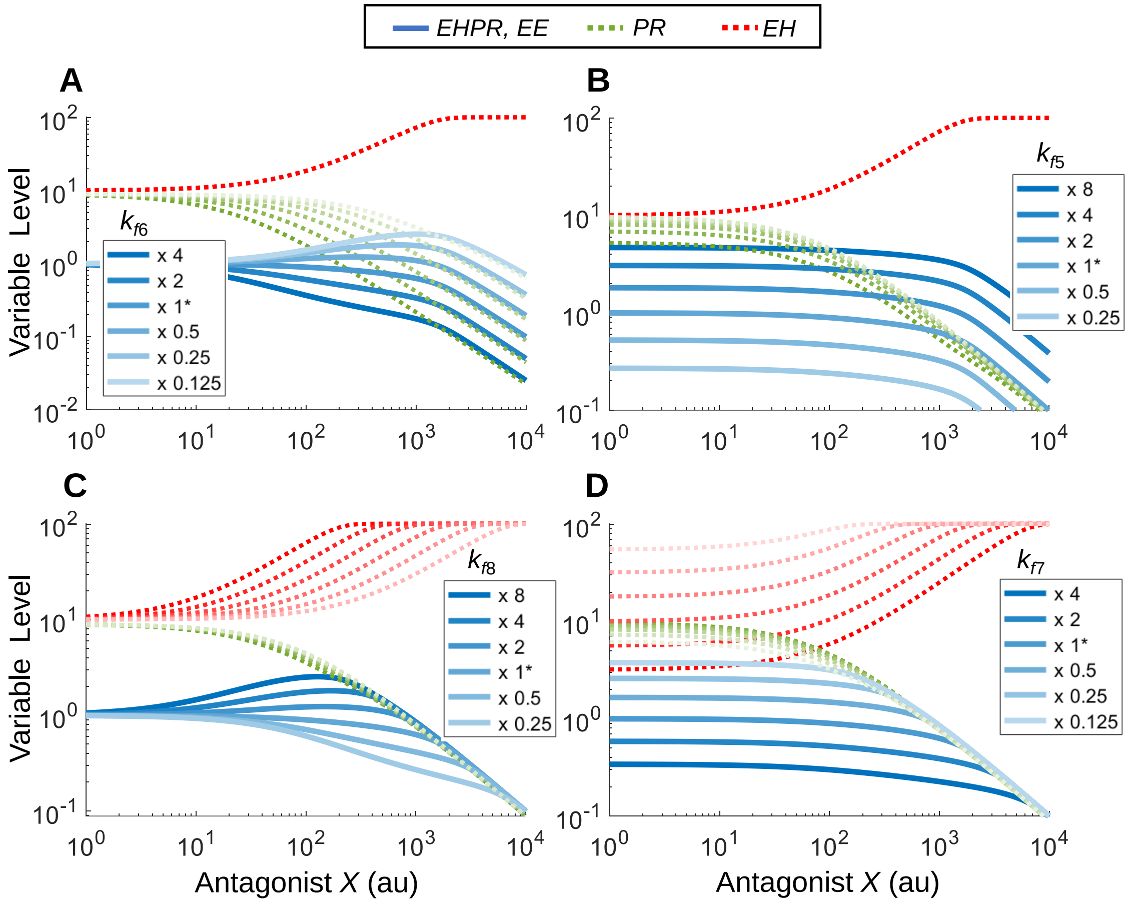


**Figure S7.** Illustration of the mechanisms of Bell-shaped DR of *EE* in response to antagonist *X* when parameters $k_{f6}$ (A), $k_{f5}$ (B). $k_{f8}$ (C), and $k_{f7}$ (D) are varied. x 1* denotes the parameter is at default value, and x 0.125, 0.25, x 0.5, x 2, x 4, and x 8 denote that the parameter is at corresponding fold of the default value. Lighter shade of the curves is associated with lower parameter values as indicated.

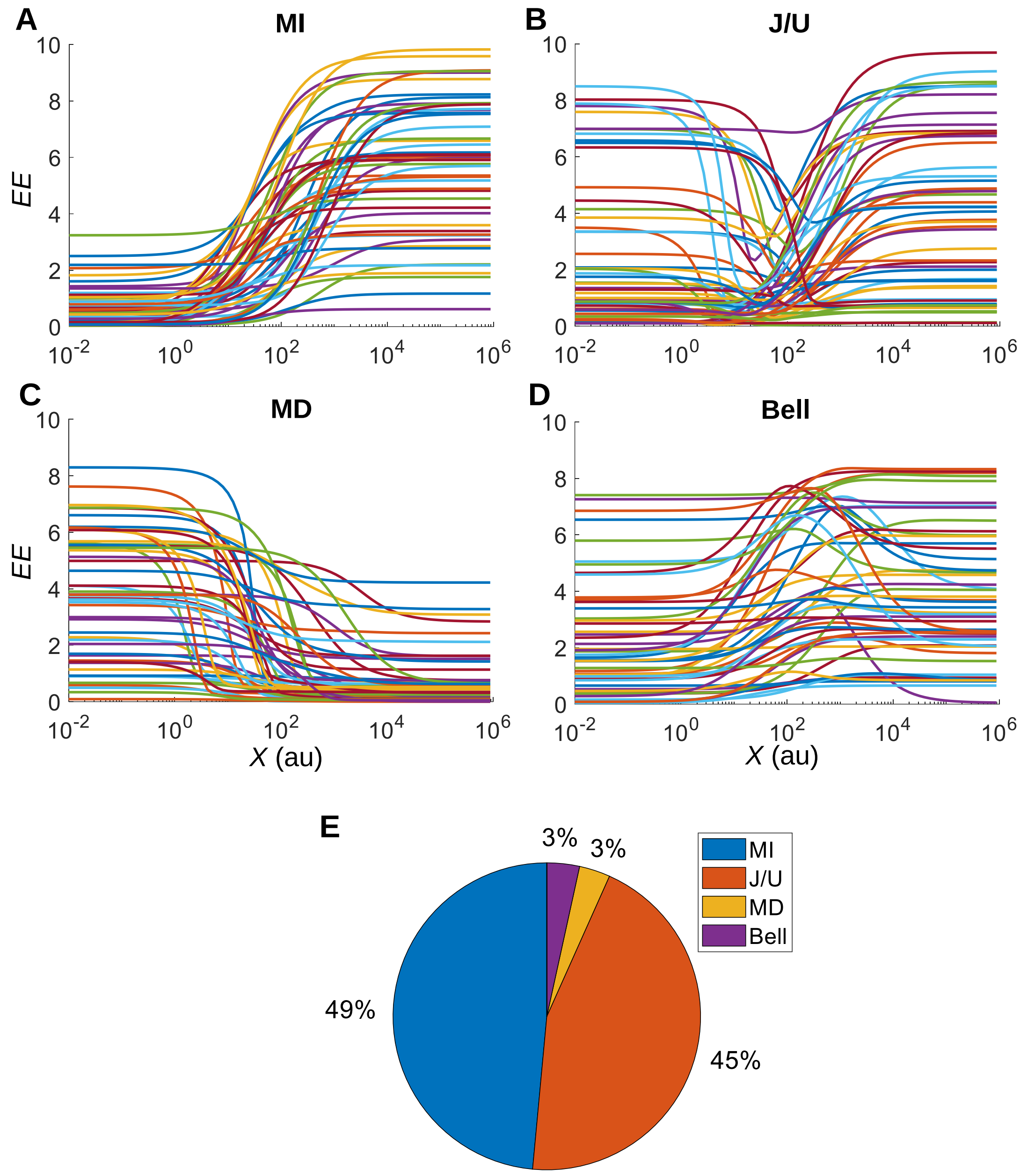


**Figure S8.** Classification of *EE* vs. *X* DR curves from 20,000 six-parameter MC simulations. The parameters $k_{f5}$, $k_{f6}$, $k_{f7}$, $k_{f8}$, were randomly sampled from uniform distributions of $\log_{10}$([0.1, 10]) as fold change relative to the respective default values, and $\omega_p$ and $\omega_c$ were randomly sampled from the uniform distribution [0,1]. **(A-D)** Randomly selected 50 DR curves for each shape as indicated. **(E)** Fractions of DR curve shapes. MI: monotonically increasing, MD: monotonically decreasing.

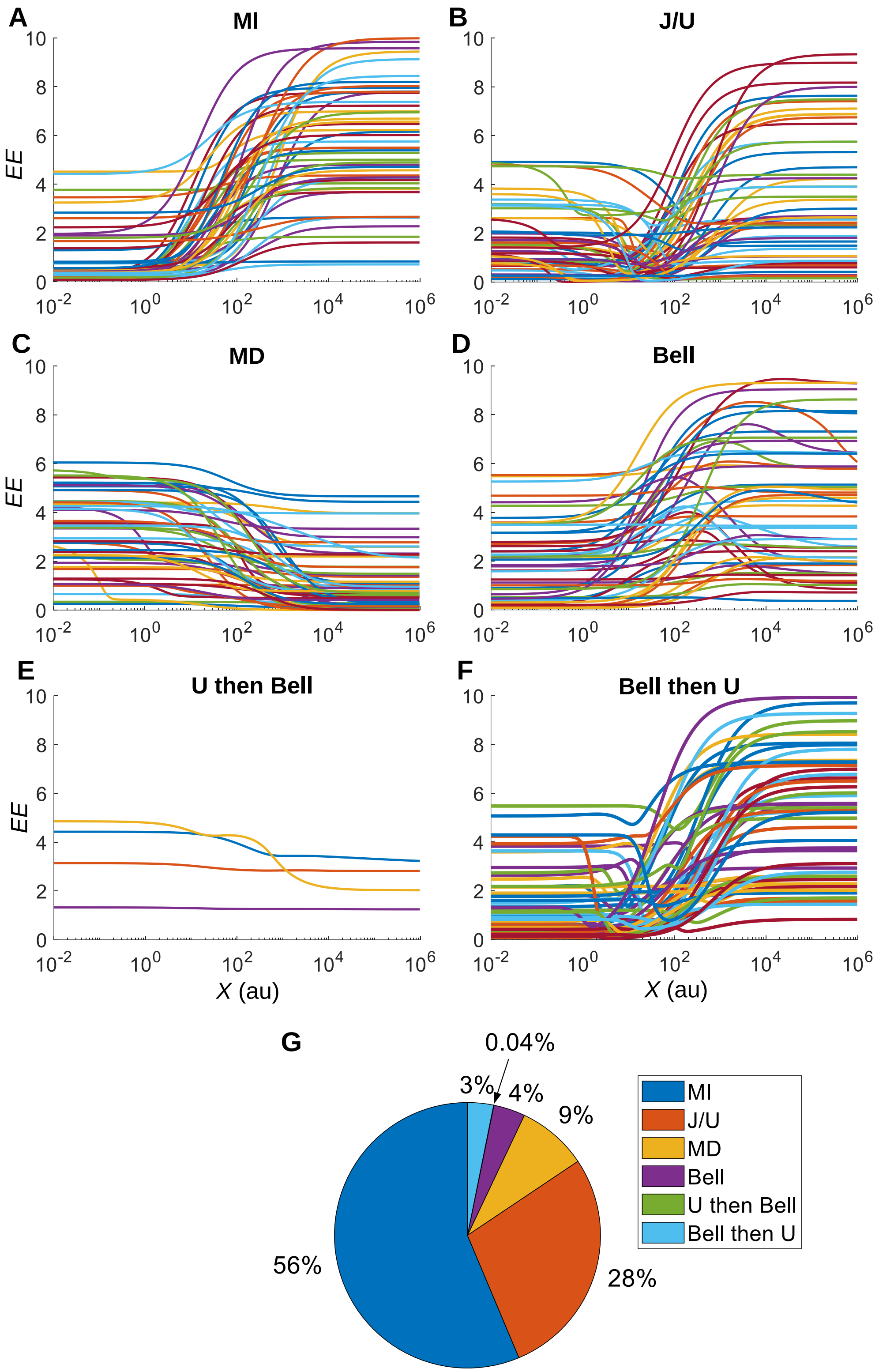


**Figure S9.** Classification of *X* vs. *EE* DR curves from MC simulations of a virtual population of 9,996 individuals. **(A-D, F)** Randomly selected 50 DR curves for MI, J/U, MD, Bell, and Bell-then-U curves respectively as indicated. **(E)** All U-then-Bell curves. Note for some curves in this category, the curvatures are too small to visualize. **(G)** Fractions of DR curve shapes. MI: monotonically increasing, MD: monotonically decreasing.

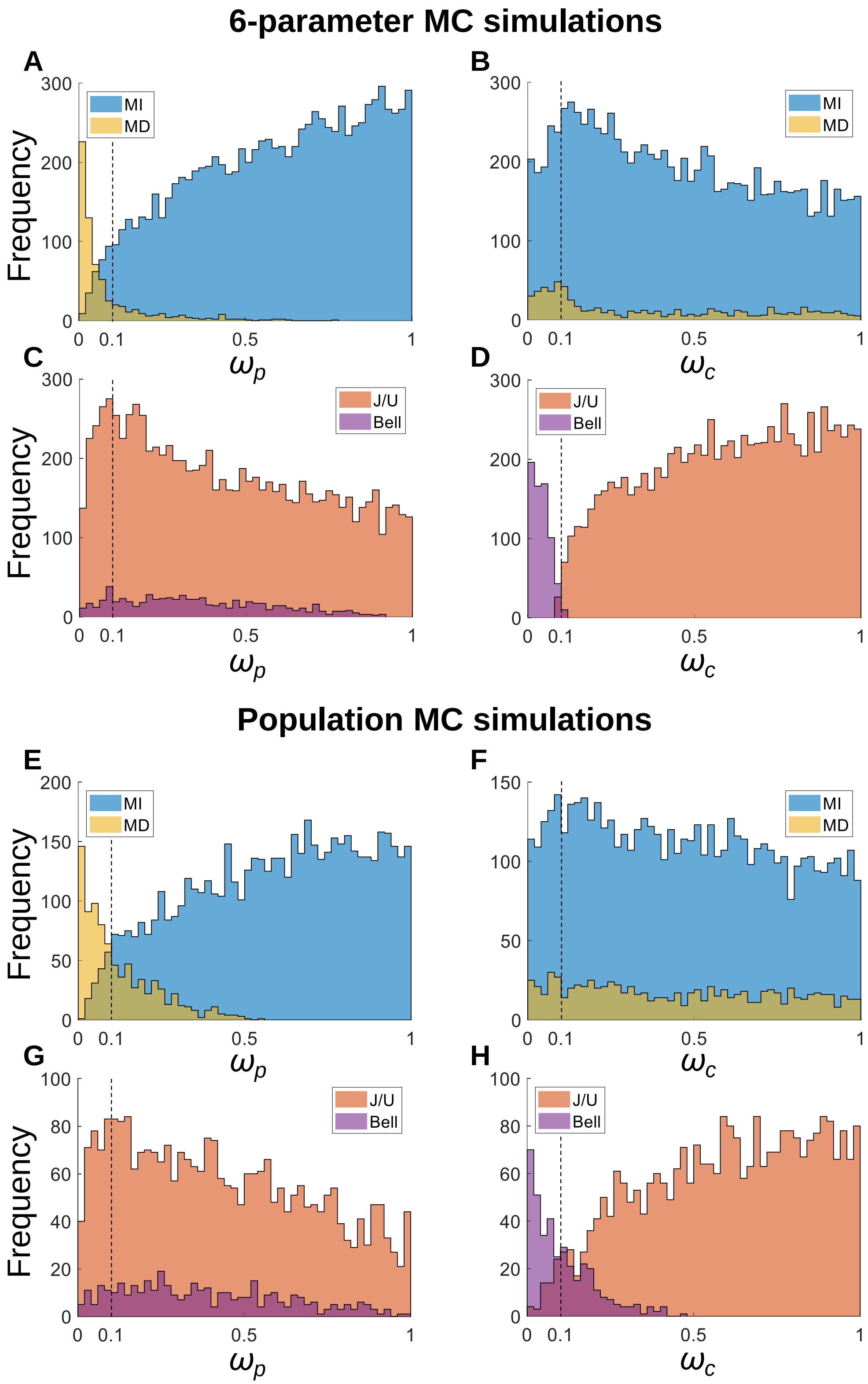


**Figure S10. Distributions of $\omega_c$ and $\omega_p$ for MI vs. MD and J/U vs. Bell curves. (A-D)** From 20,000 six-parameter MC simulations as presented in Fig.10 and S8. **(E-H)** From population MC simulations of 9,996 individuals as presented in Fig. 12 and S9. The vertical dash lines correspond to $\omega_c$ or $\omega_p$ = 0.1.
